# Viral impacts on plankton standing stocks, primary productivity, and biogeochemistry in a model ocean

**DOI:** 10.64898/2026.05.27.728272

**Authors:** Paul Frémont, Stephen J. Beckett, Daniel Muratore, David Demory, Eric Carr, Oliver Jahn, Christopher L. Follett, David Talmy, Debbie Lindell, Joshua S. Weitz, Stephanie Dutkiewicz

## Abstract

Viral lysis fuels the microbial loop by enhancing organic matter recycling (via the viral shunt) and can redirect organic matter toward export (via the viral shuttle). However, the global impact of viral infection mediated by shunt and shuttle pathways remains unclear. Here, we implemented viral infection and lysis processes in a global ocean ecosystem model, including a single phytoplankton (representing *Prochlorococcus*), virus (representing cyanophage), and nanozooplankton. Despite low but plausible levels of viral infection, high shunt efficiencies generated enhanced-productivity regions covering up to approximately one-half of the global ocean. For lower viral shunt efficiencies, the enhanced-productivity regions contracted abruptly, accompanied by steady declines in productivity. Viral-mediated increases in primary productivity reduced the extent of tropical oligotrophic regions at high shunt efficiencies, while lower efficiencies expanded oligotrophic areas. These results provide a path forward to developing predictive models of how viral infection and the fate of cellular lysates shape global ocean ecosystems.

## Introduction

Marine phytoplankton drive ocean primary production at the base of the marine food web (1), regulate major biogeochemical cycles (2), and contribute significantly to carbon export to the deep ocean (3–5). Marine primary productivity is tightly linked to the microbial loop – dissolved organic matter released through viral lysis, phytoplankton exudation and senescence, and zooplankton sloppy feeding is recycled via bacterial remineralization, enhancing nutrient and carbon availability (6). Despite the recognized importance of viruses in shaping microbial dynamics and biogeochemical fluxes (7–9), large-scale ecosystem models generally lack an explicit representation of viral lysis. Incorporating viral processes remains a key challenge for developing predictive, large-scale models of marine ecosystems (10–13). In this study, we address this gap by integrating a model of viral infection and lysis into a large-scale ecosystem framework and assess the impact of viruses on plankton stocks, productivity, and ecosystem functioning.

Phytoplankton viruses are highly abundant in the ocean with densities typically exceeding 10^8^*/L* in surface waters (8, 14–16) and can be major drivers of phytoplankton mortality (7, 8, 15–20). Viruses are the most abundant biological entities in marine environments, but represent only a small fraction of global biomass compared to cellular life (21). The ecological influence of viruses is mediated primarily through the impacts of host lysis and nutrient recycling (8). The infection of phytoplankton by viruses often leads to the release of most of the cellular content, the lysate, into the environment through cell lysis (7, 22, 23). The lysate can then fuel the microbial loop through release of dissolved organic matter, a process called the viral “shunt” (22, 24). Alternatively, viral lysis can catalyze aggregate formation of sticky lysates that contribute to the sinking particulate organic matter pool, a process called the viral “shuttle” (25, 26).

The viral shunt is hypothesized to enhance organic matter recycling, stimulate net primary production, and reduce energy transfer to higher trophic levels (10, 22, 24). At the global scale, early estimates suggest that viruses contribute approximately 20–50% of carbon recycling within the microbial loop (22, 24). In a regional modeling study, it has been suggested that (29±20%) of phytoplankton carbon losses from viral lysis are redirected into nutrient recycling in the California Current Ecosystem (27). At the scale of local ecosystems, seasonal intensification of viral lysis in the Sargasso Sea has been suggested to redirect organic matter toward rapid microbial recycling, enhancing local primary production and contributing to the formation of a pronounced subsurface oxygen maximum (9).

In addition to its influence on carbon cycling, the viral shunt is predicted to exert strong control on the marine nitrogen cycle (9, 28–33). Experimental studies have shown that degradation of *Phaeocystis pouchetii* lysates results in substantial inorganic nitrogen regeneration, stimulating the growth of heterotrophic bacteria and nanoflagellates (predators of the heterotrophic bacteria) and demonstrating efficient transfer of algal organic matter to the microbial community (30). Similarly, viral lysates derived from *Phaeocystis globosa* enhance both carbon and nitrogen assimilation by *Alteromonas* (34). Nitrogen released through viral lysis—particularly as ammonium, nitrite, and nitrate has also been proposed to accumulate within marine snow aggregates and particulate matter (29, 35). In addition, DOM release supported bacterial growth and virally regenerated N and P relieved nutrient limitation of cultured diatoms (29). Direct re-assimilation of lysate-derived nitrogen has also been demonstrated in *Synechococcus*, primarily as ammonium, with estimates indicating that nitrogen released by viral lysis of bacteria could sustain up to 71% of global primary production (32).

Quantitative models have attempted to assess the impacts of viral infection as part of multitrophic ecosystems. Consistent with empirical findings, these models show that viral lysis can in principle reshape carbon flow among eukaryotes, cyanobacteria, heterotrophic bacteria, viruses, and zooplankton (10). However, quantitative estimates of the contribution of viral lysis to organic matter recycling remain uncertain, in part, due to uncertainty in estimating *in situ* viral-induced mortality rates and the use of quantitative methods that suggest lower rates of mortality than anticipated (16, 36). Beyond bulk carbon and nutrient recycling, several studies in non-spatial, one-dimensional or local analysis of 3D frameworks have explored how host–virus interactions and environmental conditions modulate viral impacts on specific microbial clades (13, 37–43). Embedding viruses into ecosystem models requires reconciling the challenges of explaining the coexistence of viruses and zooplankton that share common prey while achieving plausible ecological outcomes. Recent modeling work suggests that host resistance to viral infection (relative to encounter rate theory) can facilitate coexistence over ecologically relevant regimes - and suggests that integrating viruses into global ocean models is increasingly feasible (13).

In this study, and building on modeling efforts described above, we integrate the mechanisms of viral lysis including both the shunt and shuttle in the Darwin ocean plankton model (44, 45). We term this virus-embedded version of the model: vDarwin (Fig. 1). Here, vDarwin is coupled to a three dimensional large-scale global circulation model (46) (Fig. 1A). Though the vDarwin model has been designed to be able to consider multiple plankton and viruses (47), in this study, as a proof of concept, we configure vDarwin to simulate one phytoplankton type (modeled on *Prochlorococcus*) infected by a single virus and preyed upon by a single zooplankton (Fig. 1B and C). We explicitly include a susceptible (*S*) and infected (*I*) host, such that total phytoplankton is the sum of both (*S* + *I*). The viral shunt and shuttle are represented by partitioning the lysate between dissolved organic matter (DOM) and sinking particulate organic matter (POM) compartments (Fig. 1D). The organic matter is remineralized to dissolved inorganic matter (e.g., nutrients). Although simplified, as we show, the inclusion of viruses is feasible and realistic, and enables model simulations to evaluate scenarios of how shifts in the fate of viral lysates can modulate plankton standing stocks, primary productivity, nutrient cycling, the range of oligotrophic regions, and global ecosystem functioning.

**Fig. 1.**
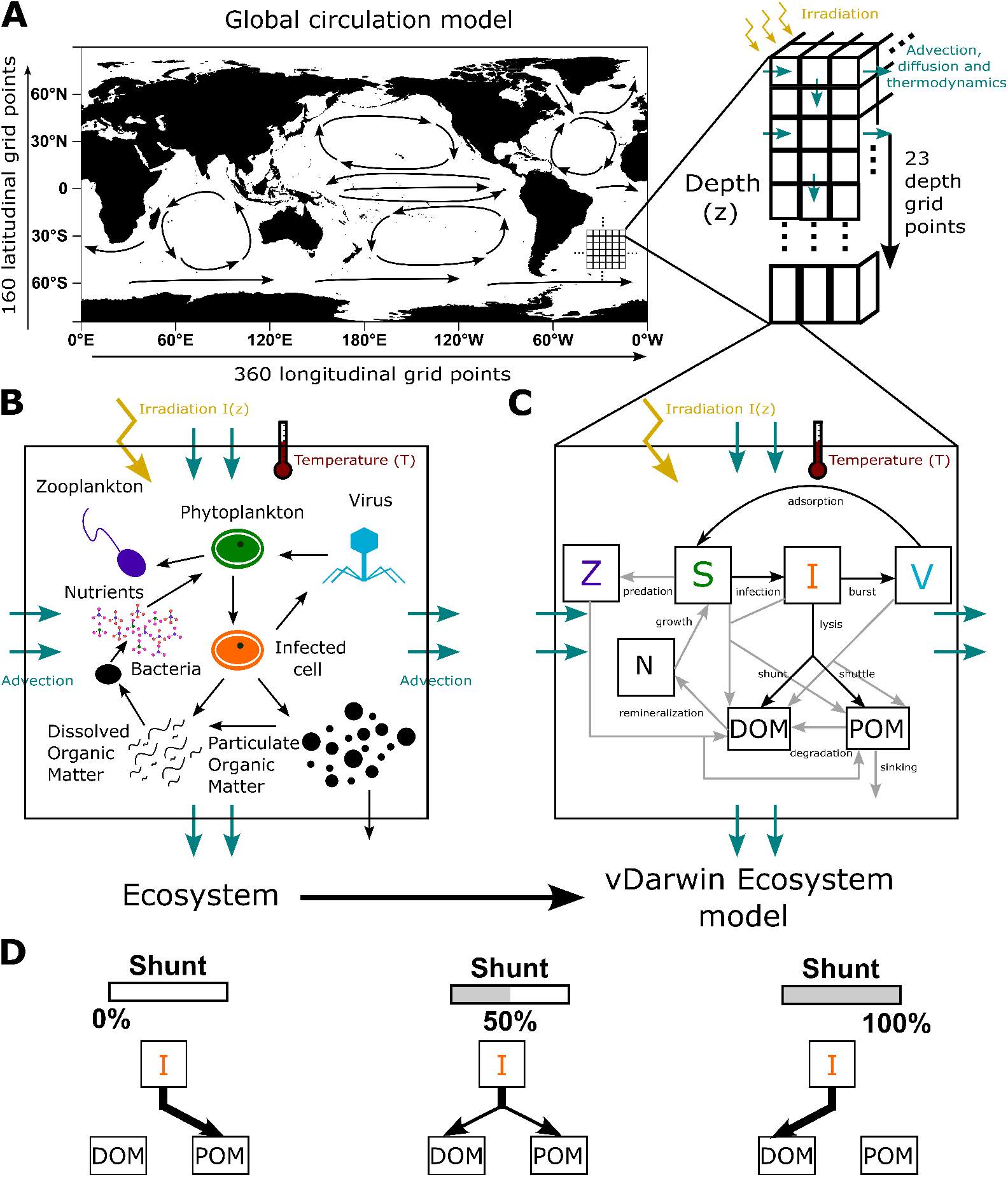
Integration of a model of viral lysis, shunt and shuttle in a large scale ocean circulation model. (**a**) Schematic of the large scale ocean circulation model (MITgcm): major surface ocean circulation patterns and the schematic of the advection/diffusion are represented. Note that vertical grid spacing increases with depth (i.e., layers become thicker). (**b**) Simplified ecosystem represented in the study. For simplicity nitrogen remineralization to nitrite and nitrate and mortality fluxes to DOM and POM are not represented. Schematic of the vDarwin ecosystem model integrated in the large scale circulation model. It represents a simplification of the ecosystem presented in (**b**) assuming that N is the limiting nutrient. It includes the growth of a single phytoplankton type, viral lysis, zooplankton grazing, the viral shunt and shuttle and remineralization (note that remineralizing bacteria are not explicitly represented). (**d**) Illustration of the viral shunt efficiency parameter representing the percentage of the viral lysate that is redirected to the DOM pool versus POM pool.

## Results

### Incorporating viral lysis into a global ecosystem model generates robust, ecologically plausible outcomes

We first assessed the qualitative behavior of the global ecosystem model, vDarwin, with idealized shunt efficiency of 100% (i.e. all of the lysate is redirected to the DOM compartment, parameter *α*, Fig. 1D, equation 2e,f, see Materials and methods). To analyze the impact of viral dynamics, we compare results of a simulation with viruses against a simulation without viruses (Fig. 2). We concentrate on results equatorward of 40^◦^N/S, since this is the region where *Prochlorococcus* is known to exist in large concentrations (48–50) (recognizing that faster growth species, not captured in this version of the model, dominate further poleward). Overall, global patterns of plankton concentrations were similar between the two simulations, though the maximal concentrations reached were lower with the explicit representation of the virus (Fig. 2A, B, D, and E). Mean total phytoplankton abundances reached count concentrations on the order of 10^8^ − 10^9^ ind.L^−1^ (~ 0.5 − 3.5 *µ*molC.L^−1^) in the most productive regions, while remaining below 10^8^ ind.L^−1^ in the core of oligotrophic regions for both simulations (Fig. 2A and B). These abundances were consistent with global estimates of *Prochlorococcus* (48–50) and regional *in situ* measurements (16, 36, 51), at least for the region of the globe (equatorward of 40^◦^N/S). Zooplankton concentrations reached concentrations on the order of 10^5^ − 10^7^ ind.L^−1^ in agreement with observations (Fig. 2D and E) (52).

**Fig. 2.**
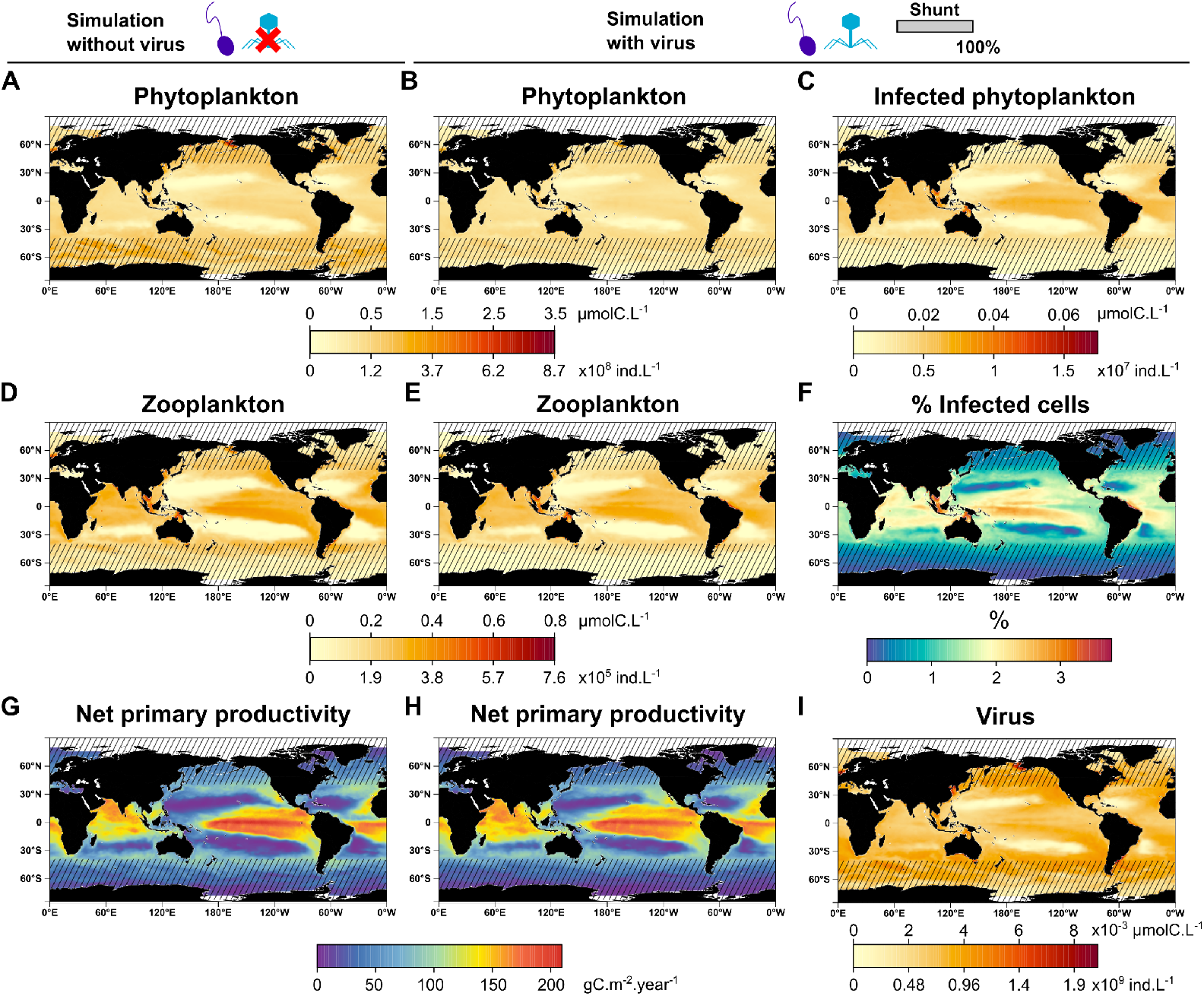
Time- and depth-averaged (tenth year, upper 100 m) model outputs for a simulation without the virus or including and explicit virus, assuming 100% viral shunt. (**a, b**) Total phytoplankton concentration from simulations without and with the virus. (**c**) Infected phytoplankton concentration. (**d, e**) Zooplankton concentration without and with the virus. (**f**) Percentage of infected cells. (**g, h**) Integrated (upper 200 m) net primary productivity without and with the virus. (**i**) Virus concentration. Striped areas indicate regions outside the main niche of *Prochlorococcus* (poleward of 40^◦^N/S).

In the simulation with explicit viral infections, the model results showed that the averaged infected-cell concentrations and the corresponding fraction of infected cells (relative to total phytoplankton concentration) were highest in the western part of the equatorial region, intermediate at mid-latitudes, and lowest within oligotrophic gyres (Fig. 2C and F). The mean fraction (first 100 m over the whole ocean) of infected cells reached a maximum of 3.8% and decreased to below 0.5% in the centers of oligotrophic regions. Notably, at Station ALOHA, the modeled mean percentage of infected cells was approximately 1%, consistent with field observations (36, 43). We further tested the sensitivity of model outputs to infected-cell representation in a model without the infected class (*SV Z* model) and in a simulation allowing for the growth of the infected class (Supplementary results, Figure S1). Finally, the model also produced realistic mean cyanophage concentrations, reaching up to ~ 2 × 10^9^ ind.L^−1^ in productive regions and ~ 5 × 10^8^ ind.L^−1^ in oligotrophic regions, with lower values in the core of gyres, in qualitative agreement with observations (16, 36) (Fig. 2I). Overall, our results confirmed that explicit parameterization of virus dynamics in a recently developed non spatial configuration (13) translated robustly to the three-dimensional vDarwin framework.

### The efficiency of the viral shunt modulates plankton standing stocks

For the phytoplankton in the simulation with 100% viral shunt efficiency, biomass was significantly higher in tropico-equatorial regions (excluding the equatorial upwelling region) in the simulation where viruses were included relative to the one without (Fig. 3A). At high latitudes (poleward of 40◦N/S), average phytoplankton biomass were strongly reduced relative to the simulation without viruses (Fig. 3A). This latter result suggests that, under the modeled viral dynamics, this phytoplankton type, does not grow fast enough to sustain high concentrations at high latitudes (note that these regions are beyond the main habitat of *Prochlorococcus*). In the tropico-equatorial regions the zooplankton also benefited strongly from viral inclusion relative to the simulation without viruses (Fig. 3D).

**Fig. 3.**
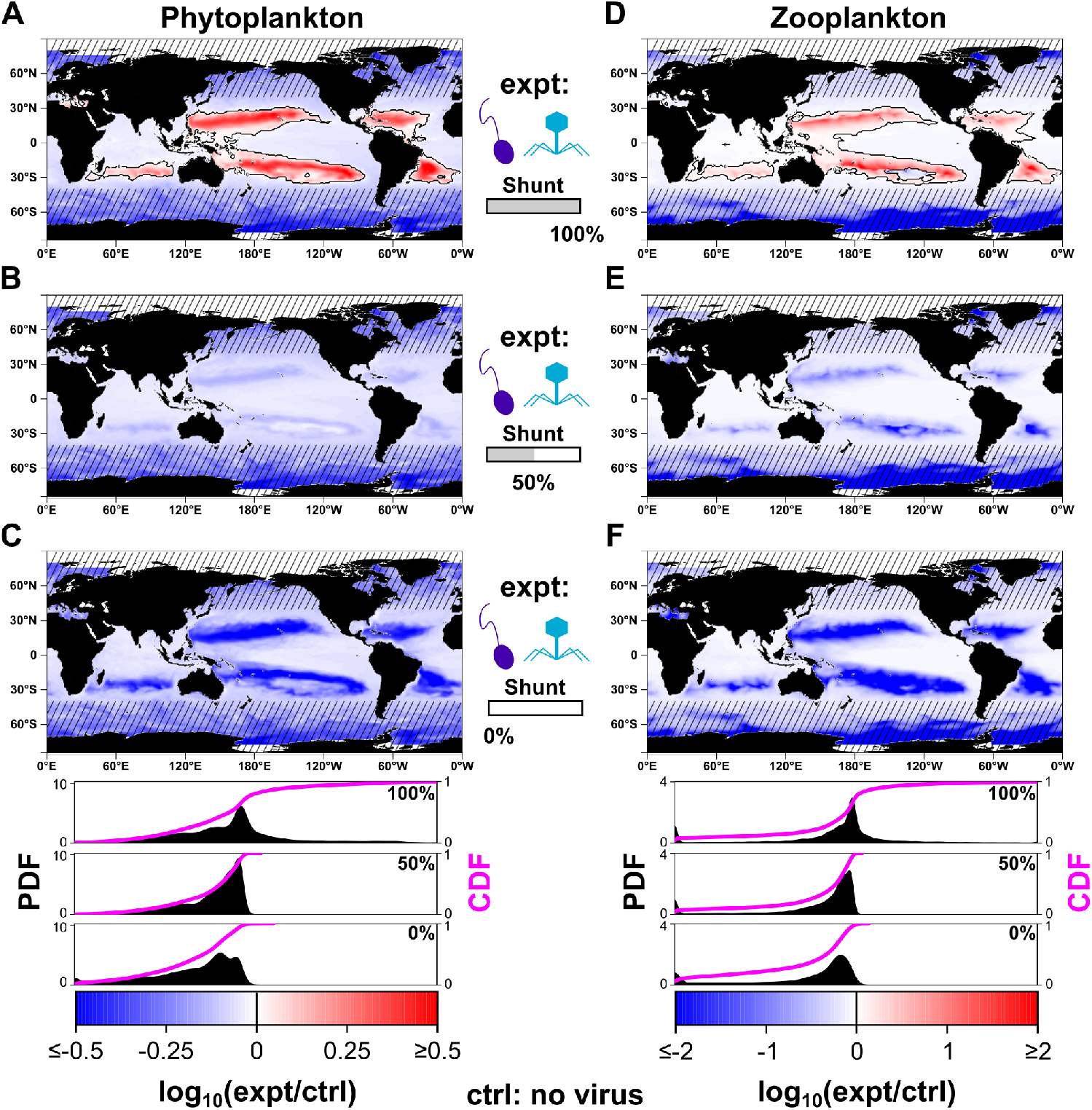
Impact of viral inclusion on plankton standing stocks in function of the viral shunt. Time- and depth-averaged (one year, upper 100 m) differences (*log*_10_ ratios) in total *Prochlorococcus* and respectively zooplankton concentration between the no virus simulation (control; ctrl) and simulation with the virus (expt) for shunt efficiencies of (**a, d**) 100, (**b, e**) 50, and (**c, f**) 0 %. Probability density function (PDF, black distribution) and cumulative distribution function (CDF, magenta) of fold changes for the different shunt efficiency values are shown in the bottom panels (data for the whole ocean). Absolute values of phytoplankton and zoo-plankton standing stocks are shown in Fig. 1A,B,D,E for the simulation without viruses and the simulation with high viral shunt efficiency. Striped areas indicate regions outside the main niche of *Prochlorococcus* (poleward of 40^◦^N/S). All log_10_ ratios are defined with the experiment (expt) simulation in the numerator and control (ctrl) simulation in the denominator. To avoid extreme values dominating the color scale, we capped absolute *log*_10_ values at the 99th percentile of the distribution or at 2 (corresponding to a 100-fold change), whichever was smaller.

To explore how these results depend on the assumption of the fate of the lysate, we conducted a series of sensitivity experiments where we varied the fraction of cellular debris going to POM versus DOM pools (Table 1, Fig. 1D). In contrast to assuming high shunt efficiency, simulations with intermediate or zero shunt efficiency (i.e. with all the lysate redirected to the POM compartment) yielded lower phytoplankton and zooplankton concentrations in tropico-equatorial regions relative to the no virus simulation (Fig. 3B, C, E, and F). Decreasing viral shunt efficiency led to lower viral concentrations in tropico-equatorial regions, while concentrations elsewhere in the ocean were slightly higher (Fig. S2). Overall, these results suggest strong effects of the virus on phytoplankton in tropico-equatorial regions, but they vary from higher to lower standing stocks depending on viral shunt efficiency. We hypothesize that higher standing stocks in the presence of the virus are likely driven by enhanced nutrient supply through remineralization resulting in increased primary productivity.

**Table 1.** Summary of simulations. “Control” refers to the reference simulation used to compute log_10_ ratio maps. The supplementary simulation “No DOM transport” refers to a simulation without advection and diffusion of dissolved organic matter (DOM).

| **Simulations** | **Virus** | **Shunt (%)** | **Other** |
| --- | --- | --- | --- |
| **Control** | No | – | – |
| **Main** | Yes | 100 | Control for supp. 1 and 2 |
|  | Yes | 90 | – |
|  | Yes | 75 | – |
|  | Yes | 60 | – |
|  | Yes | 50 | – |
|  | Yes | 25 | – |
|  | Yes | 0 | – |
| **Supplementary** | Yes | 100 | No infected type |
|  | Yes | 100 | Infected type growth |
|  | Yes | 100 | No DOM transport |

### The efficiency of the viral shunt modulates regional productivity

We assessed the effect of varying viral shunt efficiency on net primary productivity (NPP; see Materials and methods, equation 12), focusing on tropico-equatorial regions between 40◦S and 40◦N. An increase in NPP can be driven by higher turnover (increased growth rate at equal or lower standing stock), increased standing stock, or both increased growth rate and standing stock (equation 12). Absolute NPP fields for the no-virus and high-efficiency shunt simulations are shown in Fig. 2G and H. At this high viral shunt efficiency, regions of higher productivity covered approximately 50% of the ocean compared to the simulation without virus, and extending beyond regions of increased standing stocks (Fig. 4A and C). Within these regions, total annual net primary production increased by approximately 20% (Fig. 4B, red symbols and line). Spatially, NPP was higher across most tropico-equatorial regions relative to simulations without viruses, with the exception of equatorial upwelling zones (Fig. 4C).

**Fig. 4.**
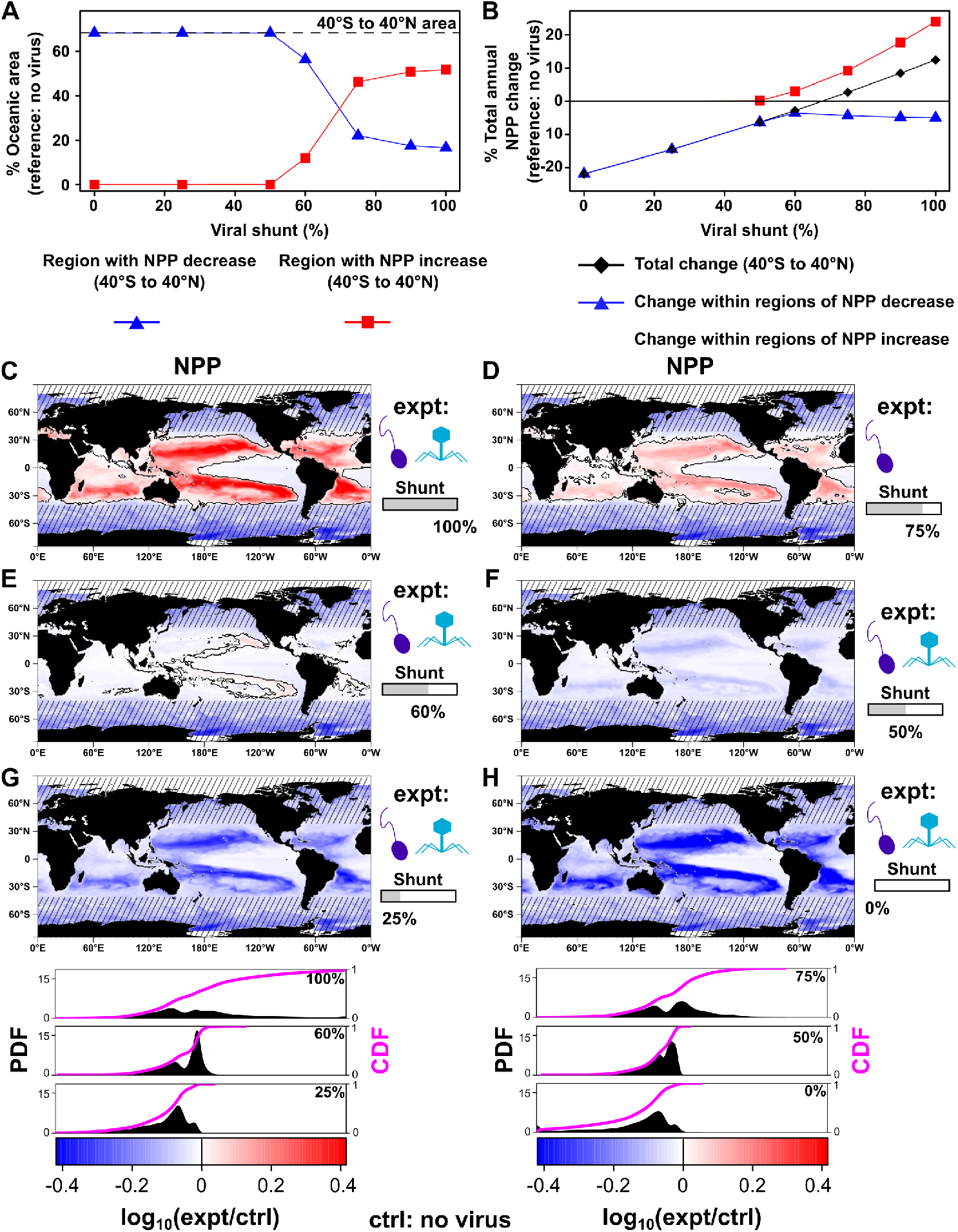
Impact of viral inclusion on primary productivity in function of the viral shunt. (**a**) Oceanic extent (percentage of total ocean area) of enhanced- and decreased-productivity regions between 40^◦^S and 40^◦^N, calculated relative to the no-virus simulation, as a function of viral shunt efficiency. The dashed line indicates the area considered (40◦S to 40◦N). (**b**) Percentage change in total annual net primary production between 40◦S and 40◦N (black), and within regions of enhanced (red) and decreased (blue) productivity. (**c**–**h**) Time-averaged and depth-integrated (one year, upper 200 m) differences (log_10_ ratios) in net primary productivity between the no-virus simulation (control; ctrl) and simulations including viruses (expt), for shunt efficiencies of (**c**) 100%, 75%, (**e**) 60%, (**f**) 50%, (**g**) 25%, and (**h**) 0%. Probability density function (PDF, black distribution) and cumulative distribution function (CDF, magenta) of fold changes for the different shunt efficiency values are shown in the bottom panels (data for the whole ocean). Absolute values of net primary productivity are shown in Fig. 1G,H for the no-virus simulation and the high viral shunt efficiency case. Striped areas indicate regions outside the main niche of *Prochlorococcus* (poleward of 40◦N/S). All log_10_ ratios are defined with the experiment (expt) simulation in the numerator and control (ctrl) simulation in the denominator. To avoid extreme values dominating the color scale, we capped absolute *log*_10_ values at the 99th percentile of the distribution or at 2 (corresponding to a 100-fold change), whichever was smaller.

The spatial extent of regions exhibiting higher productivity relative to the virus-free simulation decreased steeply with lower viral shunt efficiency (Fig. 4A,D-F). In parallel, changes in total annual net primary production declined linearly with shunt efficiency, transitioning from net positive to negative values (Fig. 4B, black dots and line). The magnitude of NPP increases also weakened progressively (Fig. 4B, D, and E), and for sufficiently low shunt efficiency, explicit viral inclusion led to negative NPP changes across the entire ocean, particularly in oligotrophic regions (Fig. 4A, B, F-H). Together, these results suggest that relatively high viral shunt efficiencies would enhance regional net primary production. The spatial extent of higher productivity in simulations including viral infection responded nonlinearly to viral shunt efficiency, declining sharply below a certain level (operationally below 75% lysate transfer to DOM in our model).

### The efficiency of the viral shunt modulates microbial loop biogeochemistry

The model considers the cycling of various biogeochemical elements, including carbon, nitrogen, and iron (see Materials and methods). In oligotrophic regions, although co-limitation exists, nitrogen is often the limiting nutrient (53, 54). Here, for simplicity, we quantified the impact of the viral shunt on nutrient cycling by examining changes (relative to the no-virus simulation) in dissolved organic nitrogen (*DON*) and dissolved inorganic nitrogen (*DIN*) concentrations (absolute values in Fig. S3A-D). Note that the model *DOM* represents only semi-labile and labile components of dissolved organic matter. We also compared the relative balance between inorganic and organic dissolved matter using the log-transformed Dissolved Nitrogen Ratio (*DNR*, see Materials and methods, equation 13). *DNR* served as a proxy for the balance between dissolved organic nitrogen accumulation and inorganic nitrogen regeneration/physical supply within the microbial loop: positive values (*DON > DIN*) indicate a relative accumulation of organic nitrogen, consistent with slower remineralization relative to dissolved organic matter production and/or physical supply and thus stronger effective nutrient limitation, whereas negative values indicate relatively greater inorganic nitrogen availability, consistent with more efficient remineralization and/or stronger external nutrient supply. As expected, positive *DNR* values occurred within oligotrophic regions where nutrients are most limiting (Fig. S3E and F).

For high shunt efficiencies, both *DON* and *DIN* were higher (relative to the no virus simulation) across regions where productivity was also higher, but with different spatial patterns (Fig. 5A and D). *DON* was broadly higher across the domain, whereas *DIN* was lower primarily at the periphery of oligotrophic regions, but to a lesser extent in their core. As a result, *DNR* was higher in the core and was lower in surrounding regions (Fig. 5G). This pattern suggests a spatial decoupling between primary production and remineralization in the core of oligotrophic regions, where the microbial loop may be less efficient due to limited viral production, which is itself limited by low primary production. This interpretation was supported by an additional simulation where *DOM* was artificially not allowed to be transported, in which core oligotrophic regions exhibited lower NPP relative to the no-virus simulation (Fig. S4). This suggests that the observed increase in NPP in oligotrophic cores was sustained by higher lateral *DON* supply from surrounding waters when viruses were present (Fig. S4). In contrast, at the periphery of oligotrophic regions, higher *DON* and *DIN*, together with lower *DNR*, were consistent with enhanced recycling within the microbial loop and a relaxation of nutrient limitation (Fig. 5A and B). Accordingly, when classifying regions of enhanced production based on changes in *DNR*, the largest absolute increases in productivity occurred where *DNR* was higher (Fig. S5A vs. S5B). In contrast, for low shunt efficiency, changes in *DON*, *DIN*, and *DNR* reversed as NPP responses became negative with the inclusion of viruses (Fig. 5B, C, E, F, H, and I). Together, these results suggest that the viral shunt might strongly affect microbial loop biogeochemistry through substantial changes in *DON*, *DIN*, and their relative balance (*DNR*). In particular, at high shunt efficiency, nutrient cycling became spatially decoupled between the core and periphery of oligotrophic regions. Note that while we projected viral impacts on nutrient cycling in the model, our microbial loop was simplified. Other features of the microbial loop that we haven’t modeled, such as phytoplankton utilization of DOM, non-labile DOM, biological nitrogen fixation, or explicit bacterial remineralization, may reduce the extent to which this ratio reflects nutrient limitation *in situ*.

**Fig. 5.**
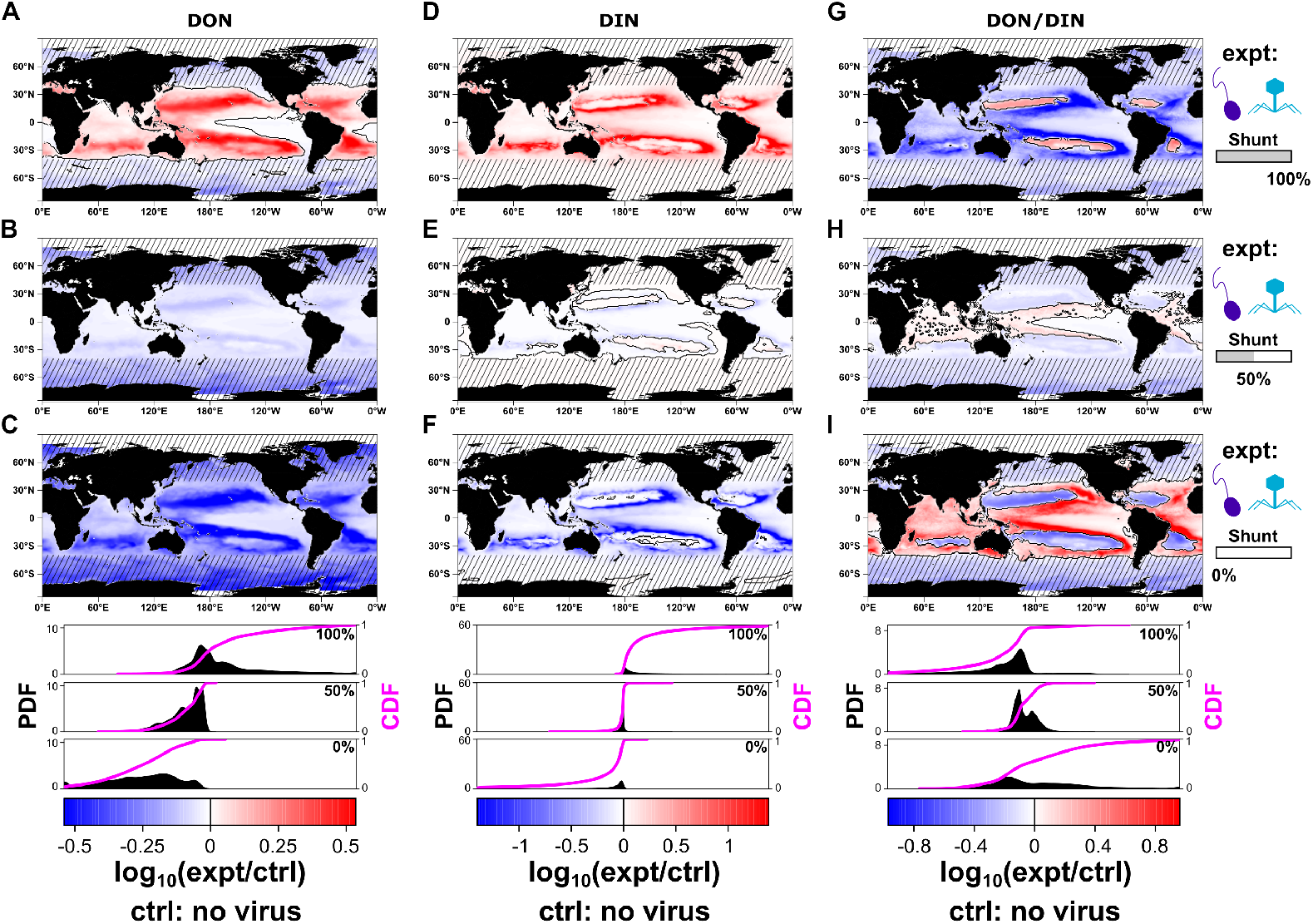
Impact of viral inclusion on *DON*, *DIN* and the *DON/DIN* ratio in function of the viral shunt. Time- and depth-averaged (one year, upper 100 m) differences in *DON*, *DIN*, and log_10_(*DON/DIN*) (hereafter *DNR*) between the no-virus simulation (control; ctrl) and simulations including viruses (expt) are shown. Panels (**a**–**c**), (**d**–**f**), and (**g**–**i**) correspond to *DON*, *DIN*, and *DNR*, respectively. Within each row, shunt efficiencies are (**a, d, g**) 100%, (**b, e, h**) 75%, and (**c, f, i**) 50%. Probability density function (PDF, black distribution) and cumulative distribution function (CDF, magenta) of fold changes for the different shunt efficiency values are shown in the bottom panels (data for the whole ocean). Absolute values of *DON*, *DIN*, and *DNR* are shown in Fig. S3A-F for the no-virus simulation and the high viral shunt efficiency case. Striped areas indicate regions outside the main niche of *Prochlorococcus* (poleward of 40◦N/S). All log_10_ ratios are defined with the experiment (expt) simulation in the numerator and control (ctrl) simulation in the denominator. To avoid extreme values dominating the color scale, we capped absolute *log*_10_ values at the 99th percentile of the distribution or at 2 (corresponding to a 100-fold change), whichever was smaller.

### The efficiency of the viral shunt modulates the extent and productivity of oligotrophic regions

Changing plankton standing stocks and primary productivity with the inclusion of viruses, as assessed in previous sections, suggests that the extent of oligotrophic regions might also be affected by the fate of the lysate. To test this in our model, we defined oligotrophic and hyper-oligotrophic regions following known criteria based on chlorophyll concentrations (see Materials and methods) (55–57). For high viral shunt efficiency, chlorophyll concentrations were substantially higher when viruses were included, and the positive region extended beyond that of higher phytoplankton standing stocks (Fig. 6A to D, Fig. S3I and J for absolute chlorophyll values on the same scale). These changes could be linked to a much lower carbon-to-chlorophyll ratio (*C/Chl*), which changes with the physiological state of phytoplankton (58) (Fig. S6). Strongest decreases in *C/Chl* occurred in regions where viral recycling enhances microbial loop efficiency and relaxed nutrient limitation, mirroring decreases in the *DNR* (Fig. 5G). Accordingly, changes in *C/Chl* were weaker in oligotrophic cores (Fig. S6, and Fig. S3G and H for absolute values of *C/Chl*). These patterns indicate a faster turnover rate and increased chlorophyll investment per unit biomass in regions of enhanced nutrient recycling when viruses were included. Consequently, regions of enhanced productivity either driven by higher fluxes or both higher biomass and growth rates (with viruses) exhibited a clear spatial structuring relative to changes in phytoplankton carbon and chlorophyll concentrations (Fig. S7). As a result of these changes, we found that the total extent of oligotrophic regions decreased by approximately ~ 23% (respectively −10% for oligotrophic and −39% for hyper-oligotrophic regions) when viruses were included and the shunt efficiency was high (Fig. 6A-D, G). The eastern boundaries of oligotrophic regions were significantly shifted westward by several hundreds of kilometers in all major oceans, most notably in the North Pacific. Finally, the contribution of oligotrophic regions (oligotrophic and hyper-oligotrophic combined) to total annual net primary production was also lower, but less substantially than the change in spatial extent (−23% spatial extent but −11% in total annual NPP, Fig. 6G and H). This response was driven by a substantial increase in net primary productivity in both regions relative to the no-virus simulation (Fig. S8). For lower viral shunt efficiencies, the extent of oligotrophic regions increased compared to the simulation without viruses (Fig. 6E, F, and G) with a linear increase in the extent of oligotrophic region as the viral shunt efficiency decreased (Fig. 6G). These results suggest that sufficiently high viral shunt efficiency can substantially decrease the spatial extent of oligotrophic regions while keeping a substantial contribution to total annual net primary production.

**Fig. 6.**
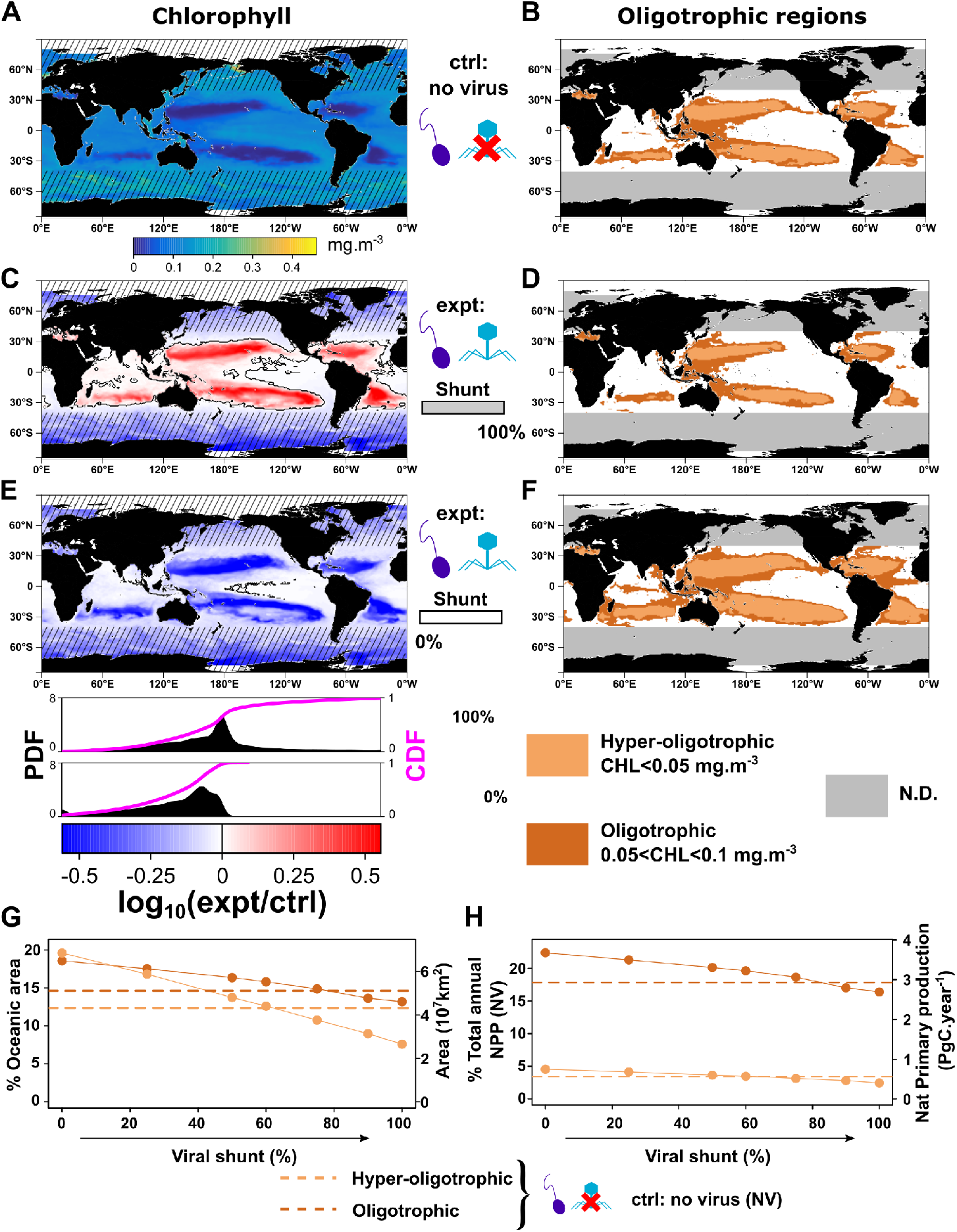
Impact of viral inclusion on chlorophyll, and the extent of oligotrophic regions in function of the viral shunt. (**a**) Time- and depth-averaged (one year, upper 100 m) modeled chlorophyll a without viruses. (**b**) Oligotrophic (brown) and hyper-oligotrophic (light brown) regions defined from mean chlorophyll thresholds for the no-virus simulation (N.D.: Not defined). (**c**) Differences (log_10_ ratios) in chlorophyll concentration between the no-virus simulation (control; ctrl) and the simulation with viruses (expt) for a shunt efficiency of 100%. (**d**) Oligotrophic (brown) and hyper-oligotrophic (light brown) regions for a viral shunt efficiency of 100%. (**e**) Same as (**c**) but for a shunt efficiency of 0%. (**f**) Same as (**d**) but for a shunt efficiency of 0%. Oligotrophic regions are not defined poleward of 40◦N/S, as the model considers a *Prochlorococcus*-only world. (**g**) Extent of oligotrophic (brown) and hyper-oligotrophic (light brown) regions as a function of viral shunt efficiency; dashed lines indicate the no-virus simulation. (**h**) Total annual net primary production within oligotrophic and hyper-oligotrophic regions and percentage relative to the no-virus simulation as a function of viral shunt efficiency; dashed lines indicate the no-virus values. Striped areas indicate regions outside the main niche of *Prochlorococcus* (poleward of 40◦N/S). All log_10_ ratios are defined with the experiment (expt) simulation in the numerator and control (ctrl) simulation in the denominator. To avoid extreme values dominating the color scale, we capped absolute *log*_10_ values at the 99th percentile of the distribution or at 2 (corresponding to a 100-fold change), whichever was smaller.

### The efficiency of the viral shunt modulates regional deep chlorophyll maxima and nutriclines

To assess how the inclusion of viruses might affect chlorophyll as well as total inorganic nutrient stocks at depth, we considered two transects, one latitudinal and one longitudinal, in the North Pacific Subtropical Gyre (NPSG, Fig. 7). For the latitudinal transect, at high viral shunt efficiency, the deep chlorophyll maximum (DCM) was most affected by the inclusion of viruses, and decreased in depth most strongly in the low-latitude part of the oligotrophic region, linked to a shoaling of the nutricline (Fig. 7A and B). Associated changes in chlorophyll concentrations and *DIN* are shown in Fig. S9. Chlorophyll was found to be higher above the DCM (with inclusion of viruses) but was lower at depth while *DIN* was higher everywhere, particularly around oligotrophic region boundaries and around and above the DCM (Fig. S9B). At low shunt efficiency, the inclusion of the virus deepened the DCM and nutricline (Fig. 7d and Fig. S9C). For the longitudinal transect, the same effects were observed mostly in the eastern part of the gyre (Fig. 7E to H). Accordingly, at high shunt efficiency, higher *DIN* was maximal in the eastern gyre with the inclusion of viruses (Fig. S9E). Subtropical gyre circulation is characterized by western intensification, with strong, narrow western boundary currents and comparatively weak flow in the eastern gyre interior (59). This circulation asymmetry likely enhances the imprint of recycling processes on nutrient structure in the eastern gyre, while more efficient advection in the west limits the accumulation of regenerated nutrients in the upper water column. These patterns suggest that under sufficiently strong viral shunt efficiency, enhanced recycling of organic matter increases nutrient availability closer to the surface contributing to the observed regional shoaling of the deep chlorophyll maximum and the shrinkage of oligotrophic regions. For the NPSG, these changes were regional and particularly affected the eastern and southern part of the gyre suggesting that the viral shunt is the most efficient in fueling the microbial loop in these regions.

**Fig. 7.**
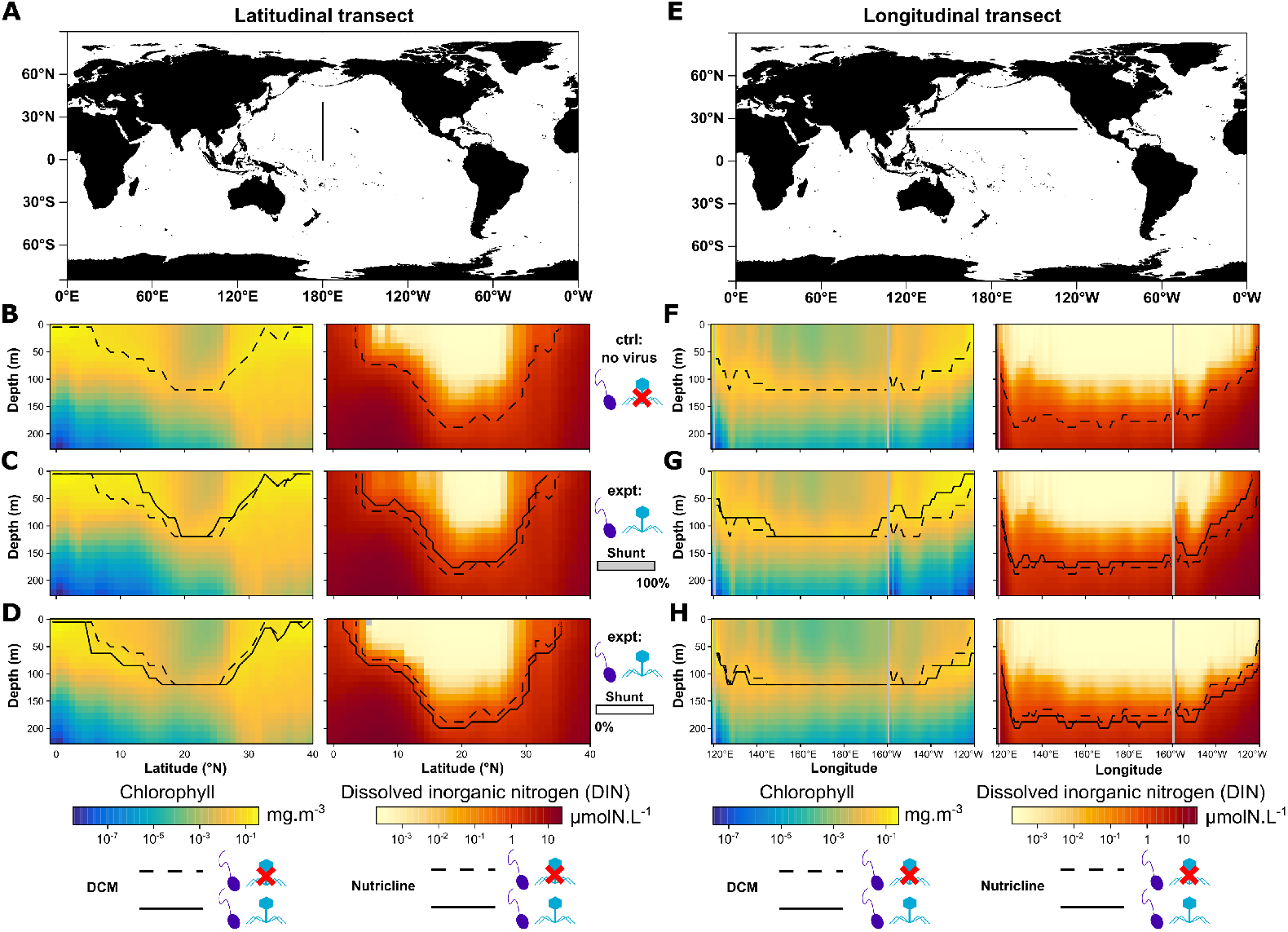
Regional changes in the deep chlorophyll maximum (DCM) and the nutricline in two transects of the North Pacific Subtropical Gyre in function of the viral shunt. (**a**) Latitudinal transect (180°E). Yearly averaged chlorophyll concentrations, DCM, dissolved inorganic nitrogen (*DIN*) and nutricline (1 *µmolN*.*L*^−1^ isocline) for (**b**) the simulation without the virus (control; ctrl) and for viral shunt efficiencies of (**c**) 100, and (**d**) 50%. (**e**) Longitudinal transect (22.5°N). Yearly averaged chlorophyll concentrations, DCM, *DIN* and nutricline for (**f**) the simulation without the virus and for viral shunt efficiencies of (**g**) 100, and (**h**) 50%. Note that concentration values are displayed on a *log*_10_ scale. For the transect data presented here, we used the approx function from R to linearly interpolate values across an equally space depth grid of 20 points (between 0 and 220 m depth). The deep chlorophyll maximum (DCM) is defined on yearly averages of chlorophyll outputs as the depth at which the average chlorophyll concentration is maximal.

## Discussion

Building on previous modeling efforts (10, 13, 19, 22, 27, 38, 43), we implemented a model of viral lysis in a global ocean ecosystem model including a representation of the viral shunt and shuttle processes. Here, we included a single phytoplankton type in both susceptible and infected cell forms, a single virus and single zooplankton grazer (though the model is designed to include multiple plankton and viruses). Qualitative evaluation of model outputs showed agreement with large scale measurements of *Prochlorococcus*, virus concentrations and the percentage of infected cells (16, 36, 43). As our model parameterized phytoplankton as a single *Prochlorococcus*, we primarily focused on viral effects in latitudes where that species is known to be abundant i.e. between 40°S and 40°N (49).

We evaluated the impact of viral inclusion relative to a simulation without viruses on global ecosystem functioning. At high viral shunt efficiencies, despite low (but plausible) levels of infection, plankton standing stocks (phytoplankton and zooplankton) increased mostly in oligotrophic gyres while primary production and chlorophyll increased across a wider region encompassing most tropical and equatorial regions, except the equatorial upwelling region (where nutrient supplies were already sufficient). This result extends and further supports projected effects of viruses on net primary production at the global scale (7, 10, 22). Previous, non-spatial models projected increases of net primary production in the presence of viruses and projected a decreased trophic transfer to nanozooplankton (10, 24). Increased trophic transfer of viral lysis products to nanozooplankton (heterotrophic nanoflagellates), mediated by enhanced bacterial grazing, has been shown experimentally (30). Here, at high shunt efficiency, we observed an increase in trophic transfer to nanozooplankton in oligotrophic regions and slight decreases in the surroundings, highlighting the importance of the global scale model and differential regional effects. These changes in standing stock, productivity and trophic transfer were also accompanied by changes in the carbon-to-chlorophyll ratio of the phytoplankton. The strongest decrease in C/Chl was in the periphery of oligotrophic regions, consistent with higher turnover rates and physiological acclimation under relaxed nutrient stress (58). This resulted in a spatial decoupling in viral effects on the nutrient cycling between the periphery of oligotrophic regions, where nutrient limitation was relaxed, and the core of these regions, where nutrient limitation remained strong. As suggested by a recent study, transport (advection and diffusion) of dissolved organic matter (DOM) can sustain primary production in oligotrophic regions (60). Our results extend the impact of this mechanism by suggesting that viral lysis–derived DOM, transported from the oligotrophic periphery, can sustain enhanced production in the oligotrophic core. Additionally, relaxation in nutrient limitation and increased chlorophyll concentrations yielded a decrease in the spatial extent of both oligotrohic and hyper-oligotrophic regions. Overall, these substantial biogeochemical changes under high shunt efficiency indicate a potentially significant role for viruses, as has been anticipated (7, 8, 10). At lower viral shunt efficiencies, the spatial extent of viral induced enhanced production was markedly lower, thereby limiting the global impact of the viral shunt in these cases. In addition, the relative increase in total annual net primary production within those regions declined linearly as viral shunt efficiency decreased. These results emphasize the need to better parameterize the balance between the viral shunt and shuttle in future modeling efforts and for further field and experimental characterization. *In situ* evidence and experimental evidence suggest that the viral shunt should increase primary production (9, 29, 32). While increased primary production due to viral infection and lysis is expected, the realized spatiotemporal extent and magnitude of this increased production remain uncertain and highly dependent on the fate of the lystate.

In this study, we focused on analyzing relative responses across simulations rather than performing direct comparisons with global-scale observations. The coarse-resolution configuration exhibits known biases in gyre extent and deep chlorophyll maximum (DCM) depth linked to unresolved mesoscale physics (61, 62). Additionally, we modeled a single phytoplankton type parameterized as a slow growing *Prochlorococcus* type. Follow-up studies are needed for a direct comparison with regional *in situ* measurements (9, 15, 16, 20, 36, 43, 63). While vDarwin included environmental controls on viral dynamics through temperature, host growth (indirectly), stratification, and advection, further developments in the viral module and microbial loop representation will enable further characterization of viral impacts, comparison with *in situ* data, and guide new observational and experimental studies.

Extensions to vDarwin could further resolve ecological and biological complexities. Viral infection and lysis rates are likely modulated by host physiological state and environmental conditions (39, 42, 64). Accounting for the diversity of phytoplankton hosts (45) and virus–host trait variation (13, 65) will further enhance ecosystem complexity and representativeness. Previous experimental and modeling studies have shown that viruses can enhance microbial coexistence and diversity (66–69). Field observations reveal complex seasonal turnover of bacteriophages and eukaryotic viruses, particularly in high-latitude systems (70). Viral infection of coccolithophores may trigger rapid bloom termination, alter carbon export pathways (71) but also follow “boom-and-busted” dynamics (40). Resolving such ecological and trait diversity represents a major next step for vDarwin and will facilitate projection of viral impacts across various ocean biomes.

Model simulations also point the way toward untested terrain: explicitly resolving environmental or host specific modulation of the partitioning between the viral shunt and the viral shuttle will facilitate comparison with observational and experimental studies. Viral lysis can enhance aggregation and particle export under certain conditions, with community composition and ecosystem structure playing a major role (4, 72, 73). In the present model framework, the viral shuttle is represented as a first order process: redirecting organic matter to a bulk POM compartment. Resolving particle size spectra and aggregation processes will further refine assessments of the global-scale balance between recycling and export mediated by viral lysis (74, 75).

Finally, expanding and refining the microbial loop representation and parameterization will allow finer-scale ecosystem dynamics to be captured. Observations indicate strong coupling between heterotrophic bacteria and viral abundances across systems (76–78). In addition, experimental studies suggest that viral lysate can be partitioned into both labile and refractory DOM pools (79). As most oceanic DOM is refractory or semi-refractory, the fate of viral lysate likely depends on bacterial community composition and metabolic pathways; conversely, the availability of such substrates may also influence bacterial community structure (80, 81). Modeling remineralizing bacteria, their viruses and DOM interactions will likely enhance our fine-scale understanding of viral impacts (82, 83).

As revealed through our global ecosystem model, the efficiency of viral lysis may play a critical role as a modulator of plankton standing stocks, primary productivity, microbial loop biogeochemistry, and oligotrophic regions. While some studies suggest rapid remineralization of viral lysate (31, 32, 84), our model results suggest a need for further *in situ* studies to better determine the rate and fate of lysate dynamics for multiple host-virus systems and across ocean biomes. Moving forward, vDarwin provides a simplified, extensible framework for integrating viral processes into global ocean ecosystem models and advancing research on the role of viruses in shaping marine ecology and biogeochemical cycles.

## Materials and methods

### Model equations

#### Advection and diffusion

Each biogeochemical tracer (described below) was advected and diffused within the three-dimensional flow field simulated by the MITgcm, which solved the hydrostatic primitive equations under the Boussinesq approximation (46). The circulation was constrained by altimetric and hydrographic observations through the ECCO-GODAE state estimates (85). Transport of each tracer followed the advection-diffusion equation (illustration in Fig. 1A):

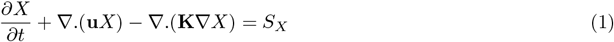

where *X* is any biogeochemical tracer, ∇ is the spatial gradient operator, **u** is the three dimensional velocity vector, **K** are the mixing coefficients and *S*_*X*_ are the source and sink fluxes of *X* (described below).

#### Ecosystem model

We developed a simplified *SIV Z* module (Susceptible-Infected-Virus-Zooplankton) integrated into the existing Darwin ecosystem model (45, 86). The Darwin model is flexible in the number of plankton it includes; here we only included a single phytoplankton (with growth parameters based on *Prochlorococcus*), but with both a susceptible *S*, and infected *I* (that does not grow by default), class, a single virus *V*, and zooplankton *Z*. The model explicitly represents other major biogeochemical tracers including chlorophyll (58, 86, 87), inorganic nutrients (ammonium, nitrite, nitrate, phosphate, iron, silica), dissolved organic matter (DOM) and particulate organic matter (POM) pools for each element (N, P, C, Fe, Si). The model also includes a representation of colored DOM (CDOM) and alkalinity. Details of equations used are described in (86). For simplicity, we only wrote model equations (in units of carbon concentration) that include the added viral processes (note that we don’t write sinking terms either, see (86)). For plankton, we have the following equations:

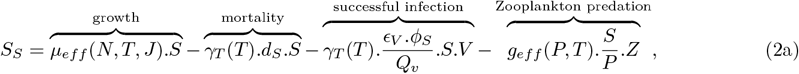

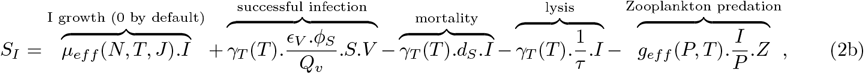

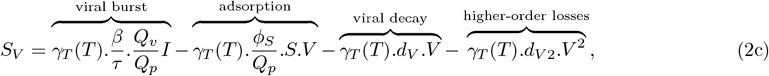

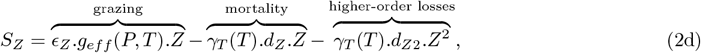

where *S*_*S*_, *S*_*I*_, *S*_*V*_ and *S*_*Z*_ are respectively the source and sink fluxes for the susceptible phytoplankton type (*S*), the infected phytoplankton type (*I*), the virus (*V*) and the zooplankton (*Z*) in (fluxes in *µmolC*.*L*^−1^.*d*^−1^ and concentrations in *µ*molC.L^−1^, or other elements). For the phytoplankton, *µ*_*eff*_ (*N, T, J*) is the effective growth rate in d^−1^ (dependent on a limiting nutrient (N), temperature (T), and light (J)), and *d*_*S*_ the mortality rate in d^−1^. For the virus, *ϵ*_*V*_ is the probability of a successful infection, *ϕ*_*S*_ is the adsorption rate of the virus in L.d^−1^, *τ* is the latent period associated to the viral infection in d, *β* is the burst size (the number of virions produced by a single lysis event), *d*_*V*_ is the viral decay rate in d^−1^, *d*_*V* 2_ is the quadratic mortality of the virus in (*µ*molC.L^−1^)^−1^.d^−1^. For the zooplankton, *g*_*eff*_ (*P, T*) is the effective grazing rate in d^−1^, *ϵ*_*Z*_ is the grazing efficiency (fraction) (88), *d*_*Z*_ the zooplankton mortality rate in d^−1^, and *d*_*Z*2_ the quadratic mortality of the zooplankton in (*µ*molC.L^−1^)^−1^.d^−1^. *Q*_*X*_ are the elemental molar quotas of each plankton type in *µ*molC.ind^−1^ (*Q*_*p*_: phytoplankton, *Q*_*v*_: virus, *Q*_*z*_: zooplankton). While *Q*_*z*_ is not used in the equations, it is used to convert zooplankton biomass to count concentrations in ind.L^−1^. *P* = *S* + *I* is the total phytoplankton concentration in *µ*molC.L^−1^, *T* is the temperature in K, *N* is the limiting nutrient concentration in *µ*molN.L^−1^ (in the case of nitrogen limitation), *J* is the irradiation in W.m^−2^, and *γ*_*T*_ (*T*) is temperature modulation that follows the Eppley curve without any temperature optimum (2, 86). While other regulations exist, we only included a temperature dependency for all virus-related processes (*γ*_*T*_ (*T*)). We list parameters values in Table S1 and further details are described in the Model parameterization section. In addition to equations of *POC* and *DOC* (and other elements) described in (86) that redistribute mortality terms to these compartments (parameters *ζ*_*X*_ in Table S1), we added the redistribution of the lysate to the *DOC* and *POC* compartments (and other elements) as described in (13):

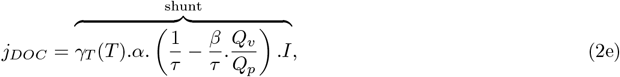

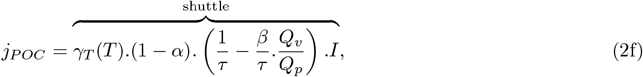

where *α* is the fraction of the lysate redirected to the *DOM* compartment. Equations are the same whatever element is considered (considering respective elemental quotas that follow the Redfield ratio (89)).

#### Chlorophyll model

Chlorophyll was computed as a function of phytoplankton biomass equation (sum of equations 2a and 2b, *P*, *S*, and *I* in mgC.L^−1^ in the following equation) and a photoacclimation factor *ρ*, representing variations in cellular Chl/C, following (58, 86, 87).

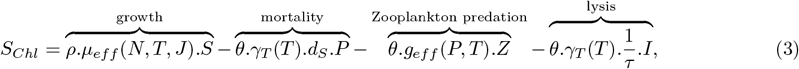

where *θ* is the chlorophyll-to-carbon ratio in gChl.gC^−1^. Reported diagnostics are converted using the inverse of *θ* (*C/Chl*), a commonly reported metric in observational studies of phytoplankton carbon-to-chlorophyll ratios (90, 91). *ρ* is the photoacclimation function following (58):

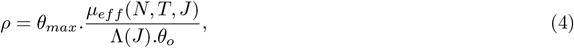

where *θ*_*max*_ is the maximum *Chl/C* ratio, *J* is the irradiance absorbed by the phytoplankton in W.m^−2^, Λ(*J*) is the light-driven photosynthetic rate in d^−1^ (see (86) for details), and *θ*_*o*_ is the acclimated *Chl/C* ratio:

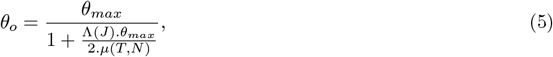

where *µ*(*T, N*) is the light-saturated photosynthesis rate as a function of nutrient and temperature only (*µ*_*S*_.*γ*_*T*_ (*T*).*γ*_*N*_ (*N*), where *µ*_*S*_ is the maximum growth rate in d^−1^). Note that in the growth term of equation 3, *S* becomes *P* in the case of the model with *I* growth.

#### Ecosystem model without the infected type

We tested an alternate representation of the virus without the explicit representation of the infected phyto-plankton type (*SV Z* model). In this case, equations of the susceptible type and zooplankton are unchanged, but the equation of the virus is as follows:

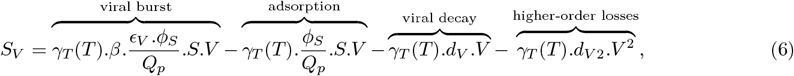

Here, virions were produced instantaneously (at a certain rate) upon encounter and successful infection of the susceptible type. Note that in this model, we have *P* = *S*.

### Model parameterization

As the basis for current model parameterization, we used the *SIV Z* model parameterization for *Prochlorococcus* and its cyanophage from (13) optimized for target concentrations of *Prochlorococcus*, its phage and nanozooplankton, in oligotrophic and mesotrophic environments. Parameter values used in the present model ocean simulations are summarized below and in Table S1.

#### Phytoplankton

We modeled a single *Prochlorococcus* phytoplankton type of radius 0.3 *µ*m that follows allometric laws for the parameterization of its maximum growth rate and half saturation constant for nutrients (13, 45). The carbon quota follows (92) and the Redfield ratio for other elements (89). The linear mortality rate of the phytoplankton is set to 10% of its maximum growth rate. The effective growth rate *µ*_*eff*_ (*N, T, J*) of the phytoplankton in d^−1^ depends on a limiting nutrient (N), temperature (T), and light (J) (86):

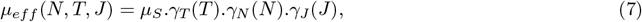

where *µ*_*S*_ is the maximum growth rate in d^−1^, *γ*_*T*_ (*T*) is the temperature dependency factor following the Eppley curve 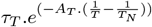, unitless) (2, 45), *γ*_N_ (*N*) is a Monod-type nutrient limitation function (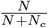, unitless) (93) for the limiting nutrient *N* following Liebig’s law of the minimum (94), *γ*_*J*_ (*J*) follows a saturating photosynthesis–irradiance curve (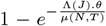, unitless) (86). Parameterization and variables used in these functions regulating phytoplankton growth are detailed in Table S2. For the default simulation, the growth rate of the infected type is 0. For the simulation with growth of the infected type, its effective growth rate follows equation 7.

#### Virus

We modeled a cyanophage of 35 nm radius, burst size *β* of 15, latent period *τ* of 0.37 d, adsorption rate *ϕ*_*S*_ of 1.7 × 10^−10^ L.d^−1^, an adsorption efficiency (the probability of successful infection) of 0.277, a linear mortality of 0.01 d^−1^ and a quadratic mortality of 23.8 (*µ*molC.L^−1^)^−1^.d^−1^ representing the viral “sweep” *lato sensu* including the consumption of viruses by higher trophic levels (95), self aggregation, and unspecific binding (96). The carbon elemental quota of the virus follows empirical elemental quota relationships for viral capsids (97). For simplicity, the stoichiometry of the virus follows the Redfield ratio (89).

#### Zooplankton

We used a Holling type II functional from for grazing (98):

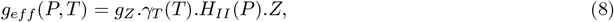

where *γ*_*T*_ (*T*) is the temperature dependency and *H*_*II*_ (*P*) is the Holling type II function:

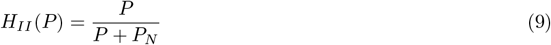

The carbon quota of the zooplankton follows (99) and the Redfield ratio for other elements (89). The linear mortality of the zooplankton is 0.067 d^−1^ and the quadratic mortality re-tuned to 0.226 (*µ*molC.L^−1^)^−1^.d^−1^ to allow for higher zooplankton concentrations. To have consistent parameters as was used in (13) (who used a Holling type I function) we set:

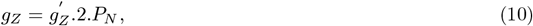

where 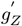 is the grazing rate used in (13) in (*µ*molN.L^−1^)^−1^.d^−1^ and *P*_*N*_ is the half saturation constant for grazing in *µ*molN.L^−1^ converted from *P*_*C*_ (Table S1) using the Redfield ratio (89). The factor 2 ensures that the grazing pressure for Holling type I (used in (13)) and Holling type II (used here) are equal for *P* = *P*_*N*_.

#### Remineralization

All remineralization rates are set to a constant of 0.033 d^−1^ (1 month^−1^) following (86). This value does not represent an explicit bacterial remineralization rate but rather an effective parameter used in the Darwin framework when heterotrophic bacteria are not explicitly resolved.

### Ecological and biogeochemical metrics

We defined the percentage of infected cells as follows:

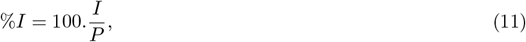

where *P* = *S* + *I* is the total phytoplankton concentration. Net primary productivity was defined as the net photosynthesis carbon production flux of the phytoplankton:

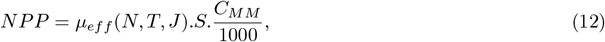

where *µ*_*eff*_ (*N, T, J*) is the effective growth rate in d^−1^ (accounting for light, temperature and nutrient limitations) and *C*_*MM*_ is the molar mass of carbon in g.mol^−1^. NPP was integrated over the upper 220 m and months yielding a unit of gC.m^−2^.y^−1^. Annual net primary production per grid cell (gC.yr^−1^) was obtained by multiplying NPP by the grid cell area (accounting for an idealized perfectly spherical Earth). Total annual net primary production over a certain domain was then obtained by summing annual net primary production over the given domain. The *log*_10_(*DON/DIN*) ratio which we termed Dissolved Nitrogen Ratio (*DNR*) was defined as follow:

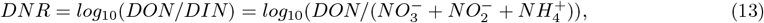

where 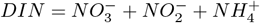 is the total dissolved inorganic nitrogen concentration (nitrate+nitrite+ ammonium) and *DON* is the dissolved organic nitrogen concentration (without counting viruses as DON). The *DNR* reflects the relative dominance of dissolved organic versus inorganic nitrogen and is used as an indicator for the balance between dissolved organic nitrogen accumulation and inorganic nitrogen regeneration/supply within the microbial loop. We used a simplified definition of the nutricline as the *DIN* isocline of 1 mmolN.m^−3^ (100). We defined oligotrophic regions as regions where time- and depth-averaged chlorophyll concentrations lay between 0.1 mg.m^−3^ and 0.05 mg.m^−3^, and hyper-oligotrophic regions as areas with concentrations below 0.05 mg.m^−3^. These thresholds are consistent with satellite-based delineations of subtropical oligotrophic regions and their hyper-oligotrophic cores (55–57).

### Simulations and visualization

Simulations were run for 10 years on a 360×160×23 spatial grid (1° resolution). Average model outputs were saved every month for all tracers. All results presented here are from the last year of simulation. To generate global average maps, all plankton and biogeochemical tracers (except NPP, see above) were averaged over the top 100 m (depth weighted averages as depth levels are unequal). For transects, tracers were averaged across months. All maps were generated using the R package maps v3.4.1 (101). For changes in all metrics, we used *log*_10_ ratios, as this provided a symmetric measure of relative change. For transect data, we used the approx function from R to linearly interpolate values across an equally spaced depth grid of 20 points (between 0 and 220 m depth).

## Acknowledgments

The authors acknowledge the University of Maryland supercomputing resources (http://hpcc.umd.edu) made available for conducting the research reported in this paper. This is a contribution of the Simons Collaboration on Ocean Processes and Ecology (SCOPE). The authors used ChatGPT to assist with coding, grammatical corrections and language clarity. The authors reviewed and edited the AI-generated content as needed and accept full responsibility for the content of the publication.

## Author Contributions

Conceptualization: PF, SJB, DM, DD, EC, OJ, CLF, DT, DL, JSW, SD; methodology: PF; investigation: PF, SD; model development: PF, SJB, DM, DD, DM, EC, OJ, CLF, DT, DL, JSW, SD; code review: EC; writing-original draft: PF; writing-review & editing: PF, SJB, DM, DD, CLF, DT, DL, JSW, SD; funding acquisition: CLF, DT, DL, JSW, SD.

## Data, code, and materials availability

All code and data necessary to generate all figures and tables are available at https://github.com/PaulFremont3/vDarwin Prochlorococcus and are archived on Zenodo at https://doi.org/10.5281/zenodo.20417458 (47). vDarwin code is available at https://github.com/jahn/darwin3. Large files (raw model outputs in .rds format) necessary to reproduce the figures are available on Zenodo (102).

## Supplementary Materials

Supplementary results

Figs. S1 to S9

Tables S1 to S2

## References

1. C. B. Field, M. J. Behrenfeld, J. T. Randerson, and P. Falkowski. “Primary production of the biosphere: integrating Terrestrial and oceanic components”. In: Science 281.5374 (1998), 237–240.

2. R. W. Eppley and B. J. Peterson. “Particulate organic matter flux and planktonic new production in the deep ocean”. In: Nature 282 (1979), 677–680.

3. S. A. Henson, R. Sanders, and E. Madsen. “Global patterns in efficiency of particulate organic carbon export and transfer to the deep ocean”. In: Global Biogeochemical Cycles 26.1 (2012).

4. L. Guidi, S. Chaffron, L. Bittner, D. Eveillard, A. Larhlimi, S. Roux, et al. “Plankton networks driving carbon export in the oligotrophic ocean”. In: Nature 532 (2015), 465–470.

5. P. Frémont, M. Gehlen, M. Vrac, J. Leconte, T. O. Delmont, P. Wincker, D. Iudicone, and O. Jaillon. “Restructuring of plankton genomic biogeography in the surface ocean under climate change”. In: Nature Climate Change 12 (2022), 393–401.

6. F. Azam, T. Fenchel, J. G. Field, J. S. Gray, L. A. Meyer-Reil, and F. Thingstad. “The ecological role of water-column microbes in the sea”. In: Marine Ecology Progress Series 10 (1983), 257–263.

7. J. A. Fuhrman and R. T. Noble. “Viruses and protists cause similar bacterial mortality in coastal sea-water”. In: Limnology and Oceanography 40.7 (1995), 1236–1242.

8. C. A. Suttle. “Marine viruses — major players in the global ecosystem”. In: Nature Reviews Microbiology 5 (2007), 801–812.

9. N. E. Gilbert, D. Muratore, C. S. Gochev, G. R. LeCleir, S. M. Cagle, H. L. Pound, C. L. Sun, A. Carrillo, K. S. Ndlovu, I. Maidanik, A. R. Coenen, L. Chittick, J. M. DeBruyn, A. Buchan, D. Lindell, M. B. Sullivan, J. S. Weitz, and S. W. Wilhelm. “Seasonal enhancement of the viral shunt catalyzes a subsurface oxygen maximum in the Sargasso Sea”. In: Nature Communications 17 (2025), 352.

10. J. S. Weitz, C. A. Stock, S. W. Wilhelm, L. Bourouiba, M. L. Coleman, A. Buchan, M. J. Follows, J. A. Fuhrman, L. F. Jover, J. T. Lennon, S. Plummer, and M. B. Sullivan. “A multitrophic model to quantify the effects of marine viruses on microbial food webs and ecosystem processes”. In: The ISME Journal 9 (2015), 1352–1364.

11. M. D. Mateus. “Bridging the gap between knowing and modeling viruses in marine systems - An upcoming frontier”. In: Frontiers in Marine Science 3 (2017).

12. D. Talmy, C. Howard-Varona, D. Eveillard, M. Covert, and M. B. Sullivan. “Viruses in multi-scale ocean models: challenges and opportunities”. In: Frontiers in Marine Science 12 (2025), 1717845.

13. P. Frémont, S. J. Beckett, D. Demory, E. Carr, C. L. Follett, D. Lindell, D. Talmy, S. Dutkiewicz, and J. S. Weitz. “Coexistence of Photosynthetic Marine Microorganisms, Viruses and Grazers: Towards Integration in Ocean Ecosystem Models”. In: Environmental Microbiology 28.4 (2026), e70295.

14. A. R. Matteson, J. M. Rowe, A. J. Ponsero, T. M. Pimentel, P. W. Boyd, and S. W. Wilhelm. “High abundances of cyanomyoviruses in marine ecosystems demonstrate ecological relevance”. In: FEMS Microbiology Ecology 84.2 (2013), 223–234.

15. K. D. Mojica, J. Huisman, S. W. Wilhelm, and C. P. Brussaard. “Latitudinal variation in virus-induced mortality of phytoplankton across the North Atlantic Ocean”. In: The ISME Journal 10 (2016), 500–513.

16. M. C. G. Carlson, F. Ribalet, I. Maidanik, et al. “Viruses affect picocyanobacterial abundance and biogeography in the North Pacific Ocean”. In: Nature Microbiology 7 (2022), 570–580.

17. Ø. Bergh, K. Y. Børsheim, G. Bratbak, and M. Heldal. “High abundance of viruses found in aquatic environments”. In: Nature 340 (1989), 467–468.

18. L. M. Proctor and J. A. Fuhrman. “Viral mortality of marine bacteria and cyanobacteria”. In: Nature 343 (1990), 60–62.

19. D. Talmy, S. J. Beckett, A. B. Zhang, D. A. A. Taniguchi, J. S. Weitz, and M. J. Follows. “Contrasting controls on microzooplankton grazing and viral infection of microbial prey”. In: Frontiers in Marine Science 6 (2019).

20. T. E. Biggs, J. Huisman, and C. P. Brussaard. “Viral lysis modifies seasonal phytoplankton dynamics and carbon flow in the Southern Ocean”. In: The ISME Journal 15 (2021), 3615–3622.

21. Y. M. Bar-On, R. Phillips, and R. Milo. “The biomass distribution on Earth”. In: Proceedings of the National Academy of Sciences 115.25 (2018), 6506–6511.

22. S. W. Wilhelm and C. A. Suttle. “Viruses and Nutrient Cycles in the Sea”. In: BioScience 49.10 (1999), 781–788.

23. J. S. Weitz. Quantitative Viral Ecology: Dynamics of Viruses and Their Microbial Hosts. Princeton, NY: Princeton University Press, 2015.

24. J. A. Fuhrman. “Marine viruses and their biogeochemical and ecological effects”. In: Nature 399 (1999), 541–548.

25. M. B. Sullivan, J. S. Weitz, and S. Wilhelm. “Viral ecology comes of age”. In: Environmental Microbiology Reports 9.1 (2017), 33–35.

26. A. E. Zimmerman, C. Howard-Varona, D. M. Needham, and et al. “Metabolic and biogeochemical consequences of viral infection in aquatic ecosystems”. In: Nature Reviews Microbiology 18 (2020), 21–34.

27. D. Talmy, S. J. Beckett, D. A. A. Taniguchi, C. P. D. Brussaard, J. S. Weitz, and M. J. Follows. “An empirical model of carbon flow through marine viruses and microzooplankton grazers”. In: Environmental Microbiology 21.6 (2019), 2171–2181.

28. M. Middelboe, N. O. G. Jørgensen, and N. Kroer. “Effects of viruses on nutrient turnover and growth efficiency of noninfected marine bacterioplankton”. In: Applied and Environmental Microbiology 62.6 (1996), 1991–1997.

29. C. J. Gobler, D. A. Hutchins, N. S. Fisher, E. M. Cosper, and S. A. Sanudo-Wilhelmy. “Release and bioavailability of C, N, P, Se, and Fe following viral lysis of a marine chrysophyte”. In: Limnology and Oceanography 42.7 (1997), 1492–1504.

30. J. Haaber and M. Middelboe. “Viral lysis of Phaeocystis pouchetii: Implications for algal population dynamics and heterotrophic C, N and P cycling”. In: The ISME Journal 3.4 (2009), 430–441.

31. E. J. Shelford, M. Middelboe, E. Friis Møller, and C. A. Suttle. “Virus-driven nitrogen cycling enhances phytoplankton growth”. In: Aquatic Microbial Ecology 66.1 (2012), 41–46.

32. E. J. Shelford and C. A. Suttle. “Virus-mediated transfer of nitrogen from heterotrophic bacteria to phytoplankton”. In: Biogeosciences 15.3 (2018), 809–819.

33. S. Wang, Y. Yang, and J. Jing. “A synthesis of viral contribution to marine nitrogen cycling”. In: Frontiers in Microbiology 13 (2022), 834581.

34. A. R. Sheik, C. P. D. Brussaard, G. Lavik, P. Lam, N. Musat, A. Krupke, S. Littmann, M. Strous, and M. M. M. Kuypers. “Responses of the coastal bacterial community to viral infection of the algae Phaeocystis globosa”. In: ISME Journal 8 (2014), 212–225.

35. A. L. Shanks and J. D. Trent. “Marine snow: microscale nutrient patches”. In: Limnology and Oceanography 24.5 (1979), 850–854.

36. N. Mruwat, M. C. G. Carlson, S. Goldin, F. Ribalet, S. Kirzner, Y. Hulata, S. J. Beckett, D. Shitrit, J. S. Weitz, E. V. Armbrust, and D. Lindell. “A single-cell polony method reveals low levels of infected Prochlorococcus in oligotrophic waters despite high cyanophage abundances”. In: The ISME Journal 15 (2021), 41–54.

37. D. Demory, L. Arsenieff, N. Simon, C. Six, F. Rigaut-Jalabert, D. Marie, P. Ge, H. Sorensen, R.-A. Sandaa, S. Jacquet, A. Sciandra, and O. Bernard. “Temperature is a key factor in Micromonas–virus interactions”. In: The ISME Journal 11 (2017), 601–612.

38. J. I. Nissimov, D. Talmy, L. Haramaty, H. F. Fredricks, E. Zelzion, B. Knowles, A. M. Eren, R. Vandzura, C. P. Laber, B. M. Schieler, C. T. Johns, K. D. More, M. J. L. Coolen, M. J. Follows, D. Bhattacharya, B. A. S. Van Mooy, and K. D. Bidle. “Biochemical diversity of glycosphingolipid biosynthesis as a driver of Coccolithovirus competitive ecology”. In: Environmental Microbiology 21.6 (2019), 2182–2197.

39. D. Demory, J. S. Weitz, A.-C. Baudoux, S. Touzeau, N. Simon, S. Rabouille, A. Sciandra, and O. Bernard. “A thermal trade-off between viral production and degradation drives virus–phytoplankton population dynamics”. In: Ecology Letters 24.8 (2021), 1773–1784.

40. K. J. Flynn, A. Mitra, W. H. Wilson, S. A. Kimmance, D. R. Clark, A. Pelusi, and L. Polimene. “‘Boom-and-busted’ dynamics of phytoplankton–virus interactions explain the paradox of the plankton”. In: New Phytologist 234.4 (2022), 1348–1360.

41. L. Xie, R. Zhang, and Y.-W. Luo. “Assessment of explicit representation of dynamic viral processes in regional marine ecological models”. In: Viruses 14.7 (2022), 1448.

42. S. Krishna, V. Peterson, L. Listmann, and J. Hinners. “Interactive effects of viral lysis and warming in a coastal ocean identified from an idealized ecosystem model”. In: Ecological Modelling 487 (2024), 110550.

43. S. J. Beckett, D. Demory, A. R. Coenen, J. R. Casey, M. Dugenne, C. L. Follett, P. Connell, M. C. G. Carlson, S. K. Hu, S. T. Wilson, D. Muratore, R. A. Rodriguez-Gonzalez, S. Peng, K. W. Becker, D. R. Mende, E. V. Armbrust, D. A. Caron, D. Lindell, A. E. White, F. Ribalet, and J. S. Weitz. “Disentangling top-down drivers of mortality underlying diel population dynamics of Prochlorococcus in the North Pacific Subtropical Gyre”. In: Nature Communications 15 (2024), 2105.

44. M. J. Follows, S. Dutkiewicz, S. Grant, and S. W. Chisholm. “Emergent biogeography of microbial communities in a model ocean”. In: Science 315 (2007), 1843–1846.

45. S. Dutkiewicz, P. Cermeno, O. Jahn, M. J. Follows, A. E. Hickman, D. A. A. Taniguchi, and B. A. Ward. “Dimensions of marine phytoplankton diversity”. In: Biogeosciences 17.3 (2020), 609–634.

46. J. Marshall, A. Adcroft, C. Hill, L. Perelman, and C. Heisey. “A finite-volume, incompressible Navier– Stokes model for studies of the ocean on parallel computers”. In: Journal of Geophysical Research: Oceans 102.C3 (1997), 5753–5766.

47. P. Frémont. PaulFremont3/vDarwin Prochlorococcus: vDarwin Prochlorococcus. Version v1. May 2026.

48. E. T. Buitenhuis, W. K. W. Li, D. Vaulot, M. W. Lomas, M. R. Landry, F. Partensky, D. M. Karl, O. Ulloa, L. Campbell, S. Jacquet, F. Lantoine, F. Chavez, D. Macías, M. Gosselin, and G. B. McManus. “Picophytoplankton biomass distribution in the global ocean”. In: Earth System Science Data 4.1 (2012), 37–46.

49. P. Flombaum, J. L. Gallegos, R. A. Gordillo, J. Rincón, L. L. Zabala, N. Jiao, D. M. Karl, W. K. W. Li, M. W. Lomas, D. Veneziano, C. S. Vera, J. A. Vrugt, and A. C. Martiny. “Present and future global distributions of the marine Cyanobacteria Prochlorococcus and Synechococcus”. In: Proceedings of the National Academy of Sciences of the United States of America 110.24 (2013), 9824–9829.

50. P. K. Lange, R. J. W. Brewin, G. Dall’Olmo, G. A. Tarran, S. Sathyendranath, M. Zubkov, and H. A. Bouman. “Scratching beneath the surface: a model to predict the vertical distribution of Prochlorococcus using remote sensing”. In: Remote Sensing 10.6 (2018).

51. R. R. Malmstrom, A. Coe, G. C. Kettler, A. C. Martiny, J. Frias-Lopez, E. R. Zinser, and S. W. Chisholm. “Temporal dynamics of Prochlorococcus ecotypes in the Atlantic and Pacific oceans”. In: The ISME Journal 4.10 (2010), 1252–1264.

52. M. Schartau, M. R. Landry, and R. A. Armstrong. “Density estimation of plankton size spectra: a reanalysis of IronEx II data”. In: Journal of Plankton Research 32.8 (2010), 1167–1184.

53. S. Dutkiewicz, B. A. Ward, F. Monteiro, and M. J. Follows. “Interconnection of nitrogen fixers and iron in the Pacific Ocean: Theory and numerical simulations”. In: Global Biogeochemical Cycles 26.1 (2012), GB1012.

54. T. J. Browning and C. M. Moore. “Global analysis of ocean phytoplankton nutrient limitation reveals high prevalence of co-limitation”. In: Nature Communications 14.1 (2023), 5014.

55. J. J. Polovina, E. A. Howell, and M. Abecassis. “Ocean’s least productive waters are expanding”. In: Geophysical Research Letters 35.3 (2008).

56. A. Morel, H. Claustre, and B. Gentili. “The most oligotrophic subtropical zones of the global ocean: similarities and differences in terms of chlorophyll and yellow substance”. In: Biogeosciences 7.10 (2010), 3139–3151.

57. J. E. O’Reilly and P. J. Werdell. “Chlorophyll algorithms for ocean color sensors - OC4, OC5 & OC6”. In: Remote Sensing of Environment 229 (2019), 32–47.

58. R. J. Geider, H. L. MacIntyre, and T. Kana. “Dynamic model of phytoplankton growth and acclimation: Responses of the balanced growth rate and the chlorophyll a:carbon ratio to light, nutrient-limitation and temperature”. In: Marine Ecology Progress Series 148.1-3 (1997), 187–200.

59. H. Stommel. “The westward intensification of wind-driven ocean currents”. In: Transactions of the American Geophysical Union 29.2 (1948), 202–206.

60. Z. Liang, R. T. Letscher, and A. N. Knapp. “Oligotrophic Ocean New Production Supported by Lateral Transport of Dissolved Organic Nutrients”. In: Global Biogeochemical Cycles 39.6 (2025). e2024.B008345 2024GB008345, e2024GB008345.

61. D. J. McGillicuddy Jr., L. A. Anderson, S. C. Doney, and M. E. Maltrud. “Eddy-driven sources and sinks of nutrients in the upper ocean: Results from a 0.1° resolution model of the North Atlantic”. In: Global Biogeochemical Cycles 17.2 (2003).

62. M. Lévy, D. Iovino, L. Resplandy, P. Klein, G. Madec, A.-M. Tréguier, S. Masson, and K. Takahashi. “Large-scale impacts of submesoscale dynamics on phytoplankton: Local and remote effects”. In: Ocean Modelling 43-44 (2012), 77–93.

63. A.-C. Baudoux, M. J. W. Veldhuis, H. J. Witte, and C. P. D. Brussaard. “Viruses as mortality agents of picophytoplankton in the deep chlorophyll maximum layer during IRONAGES III”. In: Limnology and Oceanography 52.6 (2007), 2519–2529.

64. J. A. Bonachela. “Viral plasticity facilitates host diversity in challenging environments”. In: Nature Communications 15 (2024), 7473.

65. K. F. Edwards and G. F. Steward. “Host traits drive viral life histories across phytoplankton viruses”. In: The American Naturalist 191.5 (2018), 566–581.

66. T. F. Thingstad. “Elements of a theory for the mechanisms controlling abundance, diversity, and biogeochemical role of lytic bacterial viruses in aquatic systems”. In: Limnology and Oceanography 45.6 (2000), 1320–1328.

67. S. Avrani, O. Wurtzel, I. Sharon, R. Sorek, and D. Lindell. “Genomic island variability facilitates Prochlorococcus-virus coexistence”. In: Nature 474.7353 (2011), 604–608.

68. S. Avrani, D. A. Schwartz, and D. Lindell. “Virus-host swinging party in the oceans: Incorporating biological complexity into paradigms of antagonistic coexistence”. In: Mobile Genetic Elements 2.2 (2012), 88–95.

69. J. A. Dunne, K. D. Lafferty, A. P. Dobson, R. F. Hechinger, A. M. Kuris, N. D. Martinez, J. P. McLaughlin, K. N. Mouritsen, R. Poulin, K. Reise, D. B. Stouffer, D. W. Thieltges, R. J. Williams, and C. D. Zander. “Parasites affect food web structure primarily through increased diversity and complexity”. In: PLoS Biology 11.6 (2013), e1001579.

70. G. J. Piedade, M. E. Schön, C. Lood, M. V. Fofanov, E. M. Wesdorp, T. E. G. Biggs, L. Wu, H. Bolhuis, M. G. Fischer, N. Yutin, B. E. Dutilh, and C. P. D. Brussaard. “Seasonal dynamics and diversity of Antarctic marine viruses reveal a novel viral seascape”. In: Nature Communications 15 (2024), 9192.

71. F. Vincent, M. Gralka, G. Schleyer, D. Schatz, M. Cabrera-Brufau, C. Kuhlisch, A. Sichert, S. Vidal-Melgosa, K. Mayers, N. Barak-Gavish, J. M. Flores, M. Masdeu-Navarro, J. K. Egge, A. Larsen, J.-H. Hehemann, C. Marrasé, R. Simó, O. X. Cordero, and A. Vardi. “Viral infection switches the balance between bacterial and eukaryotic recyclers of organic matter during coccolithophore blooms”. In: Nature Communications 14 (2023), 510.

72. J. S. Weitz and S. W. Wilhelm. “Ocean viruses and their effects on microbial communities and biogeochemical cycles”. In: F1000 Biology Reports 4 (2012), 17.

73. H. Kaneko, R. Blanc-Mathieu, H. Endo, S. Chaffron, T. O. Delmont, M. Gaia, N. Henry, R. Hernández-Velázquez, C. H. Nguyen, H. Mamitsuka, P. Forterre, O. Jaillon, C. de Vargas, M. B. Sullivan, C. A. Suttle, L. Guidi, and H. Ogata. “Eukaryotic virus composition can predict the efficiency of carbon export in the global ocean”. In: iScience 24.1 (2021), 102002.

74. Y. Yamada, Y. Tomaru, H. Fukuda, and T. Nagata. “Aggregate formation during the viral lysis of a marine diatom”. In: Frontiers in Marine Science 5 (2018).

75. C. P. Laber, J. E. Hunter, F. Carvalho, J. R. Collins, E. J. Hunter, B. M. Schieler, E. Boss, K. More, M. Frada, K. Thamatrakoln, C. M. Brown, L. Haramaty, J. Ossolinski, H. Fredricks, J. I. Nissimov, R. Vandzura, U. Sheyn, Y. Lehahn, R. J. Chant, A. M. Martins, M. J. L. Coolen, A. Vardi, G. R. DiTullio, B. A. S. V. Mooy, and K. D. Bidle. “Coccolithovirus facilitation of carbon export in the North Atlantic”. In: Nature Microbiology 3 (2018), 537–547.

76. C. H. Wigington, D. Sonderegger, C. P. D. Brussaard, A. Buchan, J. F. Finke, J. A. Fuhrman, J. T. Lennon, M. Middelboe, C. A. Suttle, C. Stock, W. H. Wilson, K. E. Wommack, S. W. Wilhelm, and J. S. Weitz. “Re-examination of the relationship between marine virus and microbial cell abundances”. In: Nature Microbiology 1 (2016), 15024.

77. C. Evans, J. Brandsma, M. P. Meredith, D. N. Thomas, H. J. Venables, D. W. Pond, and C. P. D. Brussaard. “Shift from Carbon Flow through the Microbial Loop to the Viral Shunt in Coastal Antarctic Waters during Austral Summer”. In: Microorganisms 9.2 (2021), 460.

78. F.-K. Shiah, C.-C. Lai, T.-Y. Chen, C.-Y. Ko, J.-H. Tai, and C.-W. Chang. “Viral shunt in the tropical oligotrophic ocean”. In: Science Advances 8 (2022), eabo2829.

79. X. Chen, W. Wei, X. Xiao, D. Wallace, C. Hu, L. Zhang, J. Batt, J. Liu, M. Gonsior, Y. Zhang, J. LaRoche, P. Hill, D. Xu, J. Wang, N. Jiao, and R. Zhang. “Heterogeneous viral contribution to dissolved organic matter processing in a long-term macrocosm experiment”. In: Environment International 158 (2022), 106950.

80. H. Osterholz, J. Niggemann, H.-A. Giebel, M. Simon, and T. Dittmar. “Inefficient microbial production of refractory dissolved organic matter in the ocean”. In: Nature Communications 6 (2015), 7422.

81. C. A. Carlson and D. A. Hansell. “Chapter 3 - DOM Sources, Sinks, Reactivity, and Budgets”. In: Biogeochemistry of Marine Dissolved Organic Matter (Second Edition). Ed. by D. A. Hansell and C. A. Carlson. Second Edition. Boston: Academic Press, 2015, pp. 65–126.

82. C. L. Follett, S. Dutkiewicz, F. Ribalet, E. Zakem, D. Caron, E. V. Armbrust, and M. J. Follows. “Trophic interactions with heterotrophic bacteria limit the range of Prochlorococcus”. In: Proceedings of the National Academy of Sciences 119.2 (2022), e2110993118.

83. R. C. Reynolds, A. C. Weiss, C. C. James, C. Y. Kojima, J. L. Weissman, J. C. Thrash, and N. M. Levine. “Defining metabolic niches for marine microbial heterotrophs”. In: Science Advances 12.17 (2026), eadz0537.

84. P. F. Hach, H. K. Marchant, A. Krupke, T. Dittmar, and M. M. M. Kuypers. “Rapid microbial diversification of dissolved organic matter in oceanic surface waters leads to carbon sequestration”. In: Scientific Reports 10 (2020), 13025.

85. C. Wunsch and P. Heimbach. “Practical global oceanic state estimation”. In: Physica D: Nonlinear Phenomena 230.1 (2007). Data Assimilation, 197–208.

86. S. Dutkiewicz, A. E. Hickman, O. Jahn, W. W. Gregg, C. B. Mouw, and M. J. Follows. “Capturing optically important constituents and properties in a marine biogeochemical and ecosystem model”. In: Biogeosciences 12 (2015), 4447–4481.

87. R. J. Geider, H. L. MacIntyre, and T. M. Kana. “A dynamic regulatory model of photoacclimation to light, nutrient and temperature”. In: Limnology and Oceanography 43 (1998), 679–694.

88. D. Straile. “Gross growth efficiencies of protozoan and metazoan zooplankton and their dependence on food concentration, predator-prey weight ratio, and taxonomic group”. In: Limnology and Oceanography 42.6 (1997), 1375–1385.

89. A. C. Redfield. “On the proportions of organic derivatives in sea water and their relation to the composition of plankton”. In: James Johnstone Memorial Volume (1934), 176–192.

90. H. H. Jakobsen and S. Markager. “Carbon-to-chlorophyll ratio for phytoplankton in temperate coastal waters: Seasonal patterns and relationship to nutrients”. In: Limnology and Oceanography 61.5 (2016), 1853–1868.

91. T. J. Smyth, G. H. Tilstone, and S. B. Groom. “Global patterns of phytoplankton carbon-to-chlorophyll ratios from ocean colour observations”. In: Frontiers in Marine Science 10 (2023), 1191216.

92. P. G. Verity, C. Y. Robertson, C. R. Tronzo, M. G. Andrews, J. R. Nelson, and M. E. Sieracki. “Relationships between cell volume and the carbon and nitrogen content of marine photosynthetic nanoplankton”. In: Limnology and Oceanography 37.7 (1992), 1434–1446.

93. J. Monod. “The growth of bacterial cultures”. In: Annual Review of Microbiology 3.1 (1949), 371–394.

94. J. v. Liebig. Die organische Chemie in ihrer Anwendung auf Agricultur und Physiologie. Braunschweig: Friedrich Vieweg und Sohn, 1840.

95. K. M. J. Mayers, C. Kuhlisch, J. T. R. Basso, M. R. Saltvedt, A. Buchan, and R.-A. Sandaa. “Grazing on Marine Viruses and Its Biogeochemical Implications”. In: mBio 14.1 (2023), e01921–21.

96. K. D. Mojica and C. P. D. Brussaard. “Marine viruses and their role in marine ecosystems and carbon cycling”. In: Annual Review of Marine Science 18.1 (2026), 351–380.

97. L. F. Jover, T. Chad Effler, A. Buchan, S. W. Wilhelm, and J. S. Weitz. “The elemental composition of virus particles: implications for marine biogeochemical cycles”. In: Nature Reviews Microbiology 12 (2014), 519–528.

98. C. S. Holling. “Some characteristics of simple types of predation and parasitism”. In: The Canadian Entomologist 91.7 (1959), 385–398.

99. S. Menden-Deuer and E. J. Lessard. “Carbon to volume relationships for dinoflagellates, diatoms, and other protist plankton”. In: Limnology and Oceanography 45.3 (2000), 569–579.

100. C. A. Garcia, S. E. Baer, N. S. Garcia, S. Rauschenberg, B. S. Twining, M. W. Lomas, and A. C. Martiny. “Nutrient supply controls particulate elemental concentrations and ratios in the low latitude eastern Indian Ocean”. In: Nature Communications 9 (2018), 4868.

101. R. A. Becker, A. R. Wilks, R. Brownrigg, T. P. Minka, and A. Deckmyn. maps: Draw Geographical Maps. R package version 3.5.0. Comprehensive R Archive Network (CRAN), 2021.

102. P. Frémont. vDARWIN Prochlorococcus model outputs. Version 1. Zenodo, Apr. 2026.

