## Supplementary Information for "Viral impacts on plankton standing stocks, primary productivity, and biogeochemistry in a model ocean"

### Supplementary results

#### Sensitivity to infected-cell representation

To assess the sensitivity of model outcomes to the explicit representation of infected cells, we performed two additional simulations: (i) a reduced $SVZ$ configuration without an infected compartment (equation 6), and (ii) a configuration allowing infected cells to continue growing prior to lysis.

Relative to the default virus simulation, the $SVZ$ configuration increased phytoplankton and zooplankton biomass in oligotrophic regions and decreased them elsewhere (Fig. S1A and C). Allowing growth of infected cells prior to lysis dampened these differences (Fig. S1B and D). Net primary production showed a similar pattern, with a stronger increase in the $SVZ$ configuration and a more dampened increase when infected cells were allowed to grow (Fig. S1G and H). For the virus, removing the infected compartment substantially increased mean stocks, primarily in core tropico-equatorial regions, whereas allowing infected-cell growth produced a weaker increase (Fig. S1E and F). Together, these experiments indicate that explicit infected-cell dynamics modulate both host and zooplankton biomass distributions and viral standing stocks.

| Parameter | Symbol | Unit | Value or Range | Reference |
|---|---|---|---|---|
| | | *Prochlorococcus* | | |
| *Prochlorococcus* maximum growth rate | $\mu_S$ | $d^{-1}$ | 0.75 | (45) |
| Infected *Prochlorococcus* maximum growth rate | $\mu_I$ | $d^{-1}$ | 0 (default) or 0.75 | (45) |
| Linear mortality of the phytoplankton | $d_S$ | $d^{-1}$ | 0.075 | Tuned |
| Phytoplankton mortality exported fraction | $1 - \zeta_P$ | fraction | 0.33 | (45) |
| Radius | $r_P$ | $\mu$m | 0.3 | Same as (13) |
| Carbon quota | $Q_p$ | $\mu$molC.ind$^{-1}$ | $4.04 \times 10^{-9}$ | (99) |
| | | *Virus* | | |
| Adsorption rate | $\phi_S$ | L.d$^{-1}$ | $1.7 \times 10^{-10}$ | (13) |
| Adsorption efficiency | $\epsilon_V$ | fraction | 0.277 | (13) |
| Linear mortality of the virus | $d_V$ | $d^{-1}$ | 0.01 | (13) |
| Quadratic mortality of the virus | $d_{V2}$ | ($\mu$molC.L$^{-1}$)$^{-1}$.d$^{-1}$ | 23.8 | (13) |
| Latent period | $\tau$ | d | 0.37 | (13) |
| Shunt fraction | $\alpha$ | fraction | $[0 - 1]$ | - |
| Viral mortality exported fraction | $1 - \zeta_V$ | fraction | 0 | - |
| Radius | $r_V$ | nm | 35 | Same as (13) |
| Carbon quota | $Q_v$ | $\mu$molC.ind$^{-1}$ | $4.18 \times 10^{-12}$ | (97) |
| | | *Zooplankton* | | |
| Grazing rate | $g_Z$ | $d^{-1}$ | 4.4 | Adapted from (13) |
| Grazing efficiency | $\epsilon_Z$ | fraction | 0.3 | (88) |
| Linear mortality of the zooplankton | $d_Z$ | $d^{-1}$ | 0.067 | (45) |
| Quadratic mortality of the zooplankton | $d_{Z2}$ | ($\mu$molC.L$^{-1}$)$^{-1}$.d$^{-1}$ | 0.226 | Tuned |
| Zooplankton mortality exported fraction | $1 - \zeta_Z$ | fraction | 0.33 | - |
| Radius | $r_V$ | $\mu$m | 2.5 | Same as (13) |
| Carbon quota | $Q_z$ | $\mu$molC.ind$^{-1}$ | $1.06 \times 10^{-6}$ | (99) |
| Half saturation constant for grazing | $P_C$ | $\mu$molC.L$^{-1}$ | 1.5 | (45) |
| | | *Remineralization* | | |
| Remineralization rate | $\kappa$ | $d^{-1}$ | 0.033 | Tuned |

**Table S1. Model parameters.**

| Regulation | Equation | Parameter or variable | Symbol | Unit | Value | References |
|---|---|---|---|---|---|---|
| Nutrient | $\dfrac{N}{N+N_c}$ | Limiting nutrient concentration | $N$ | $\mu$mol.L$^{-1}$ | - | (94) |
| | | Half-saturation constant | $N_c$ | $\mu$mol.L$^{-1}$ | Allometric laws (45) | (93) |
| Temperature | $\tau_T\, e^{-A_T\left(\frac{1}{T}-\frac{1}{T_N}\right)}$ | Scaling factor | $\tau_T$ | - | 0.8 | |
| | | Temperature sensitivity | $A_T$ | K | -4000 | (2) |
| | | Temperature | $T$ | K | - | (45) |
| | | Reference temperature | $T_N$ | K | 293.15 | |
| Light | $1-e^{-\frac{\Lambda(J)\,\theta}{\mu(N,T)}}$ | Light-driven photosynthetic rate | $\Lambda(J)$ | d$^{-1}$ | - | |
| | | Chl/C ratio | $\theta$ | mgChl.mgC$^{-1}$ | - | (86) |
| | | Light-saturated growth rate | $\mu(N,T)$ | d$^{-1}$ | - | |

**Table S2. Functional forms of phytoplankton growth regulation and parameterization.** The limiting nutrient follows Liebig's law of the minimum (94) and can either be iron, a form of nitrogen (ammonium, nitrate or nitrite) or phosphorous. Details of the computation of the light-driven photosynthetic rate is given in (86).

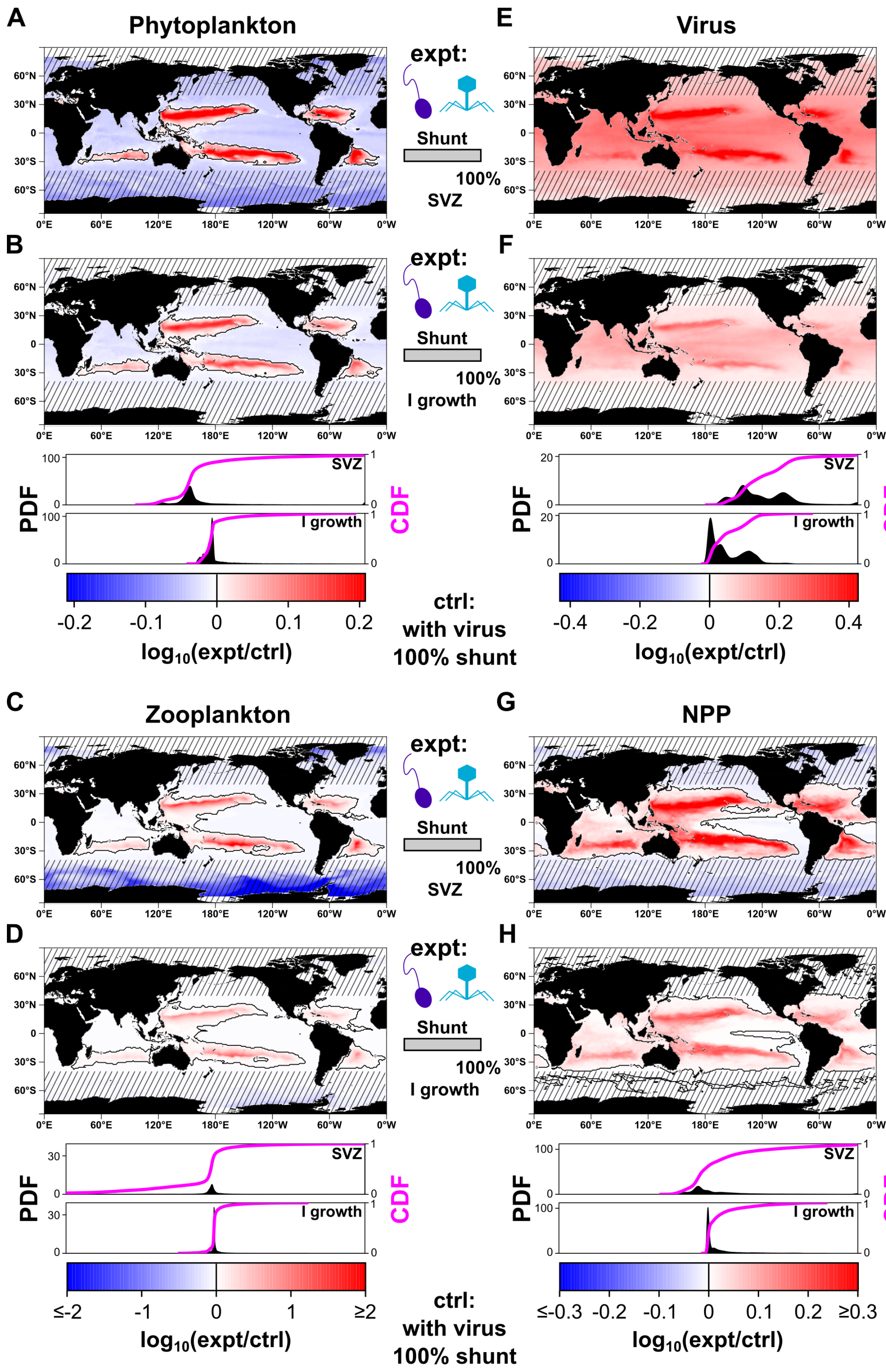


**Fig. S1. Impact of removing the infected type or allowing for growth of the infected type on plankton standing stocks and net primary productivity.** Fold changes ($log_{10}$ ratios) in time- and depth-averaged (one year, upper 100 m) model outputs between the simulation assuming 100% viral shunt efficiency (control; ctrl) and simulations respectively without the infected type and with the infected type for (**a**, **b**) phytoplankton concentrations, (**c**, **d**) zooplankton concentrations (**e**, **f**) virus concentrations, and (**g**, **h**) net primary productivity (NPP). Probability density function (PDF) and cumulative distribution function (CDF) of fold changes are shown in the bottom panels (data for the whole ocean). Striped areas indicate regions outside the main niche of *Prochlorococcus* (poleward of 40°N/S). All $\log_{10}$ ratios are defined with the experiment (expt) simulation in the numerator and control (ctrl) simulation in the denominator.

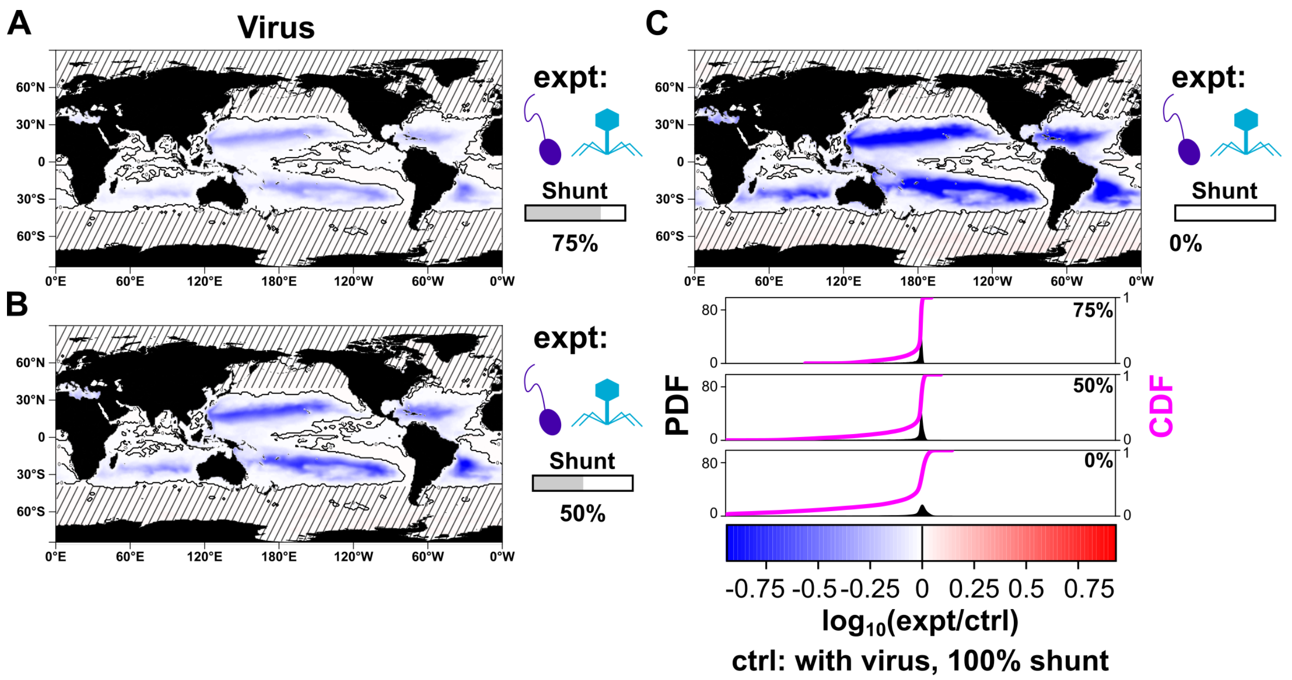


**Fig. S2. Impact of the viral shunt efficiency on the viruses concentrations.** Fold changes ($log_{10}$ ratios) in time- and depth-averaged (one year, upper 100 m) model viruses concentration between the simulation assuming 100% viral shunt efficiency (control; ctrl) and simulations with (**a**) 75, (**b**) 50, and (**c**) 0% shunt efficiency. Probability density function (PDF, black distribution) and cumulative distribution function (CDF, magenta) of fold changes for the different shunt efficiency values are shown in the bottom panels (data for the whole ocean). Striped areas indicate regions outside the main niche of *Prochlorococcus* (poleward of 40°N/S). All $\log_{10}$ ratios are defined with the experiment (expt) simulation in the numerator and control (ctrl) simulation in the denominator. To avoid extreme values dominating the color scale, we capped absolute $log_{10}$ values at the 99th percentile of the distribution or at 2 (corresponding to a 100-fold change), whichever was smaller.

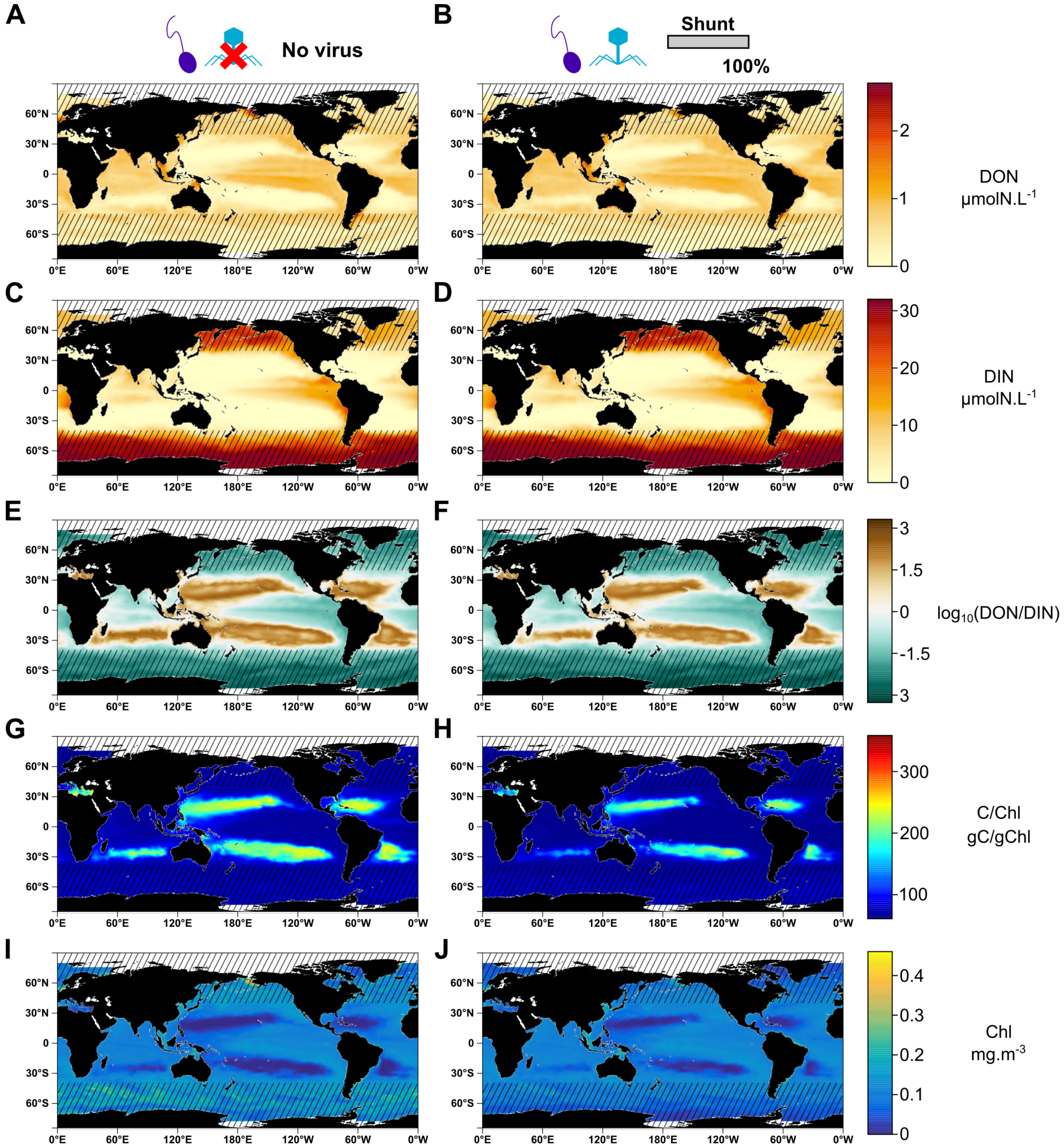


**Fig. S3. Average maps of various biogeochemical tracers for the simulation with and without the virus assuming a high viral shunt efficiency.** Yearly averages of biogeochemical tracers respectively for the simulations without the virus and with the virus: (**a**, **b**) $log_{10}(DON/DIN)$ (termed $DNR$, $DON$: dissolved organic nitrogen and $DIN$: dissolved inorganic nitrogen), (**c**, **d**) carbon-to-chlorophyll ratio ($C/Chl$) and (**e**, **f**) chlorophyll (the chlorophyll map for the simulation without the virus is also shown in Fig. 6A). They are reproduced here to facilitate visual comparison under a consistent color scale. Striped areas indicate regions outside the main niche of *Prochlorococcus* (poleward of 40°N/S).

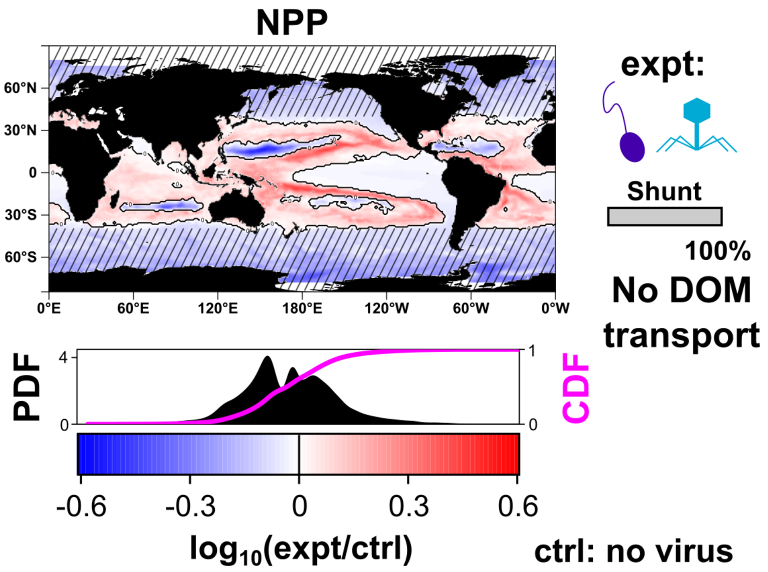


**Fig. S4. Impact of viral inclusion without** ***DOM*** **transport on changes in net primary productivity.** Fold changes ($log_{10}$ ratios) in time-averaged and depth-integrated (one year, upper 200 m) model net primary productivity between the simulation assuming 100% viral shunt efficiency without $DOM$ transport and the simulation without the virus (with $DOM$ transport, control; ctrl). Striped areas indicate regions outside the main niche of *Prochlorococcus* (poleward of 40°N/S). All $\log_{10}$ ratios are defined with the experiment (expt) simulation in the numerator and control (ctrl) simulation in the denominator. To avoid extreme values dominating the color scale, we capped absolute $log_{10}$ values at the 99th percentile of the distribution or at 2 (corresponding to a 100-fold change), whichever was smaller. Probability density function (PDF, black distribution) and cumulative distribution function (CDF, magenta) of fold changes are shown in the bottom panels (data for the whole ocean).

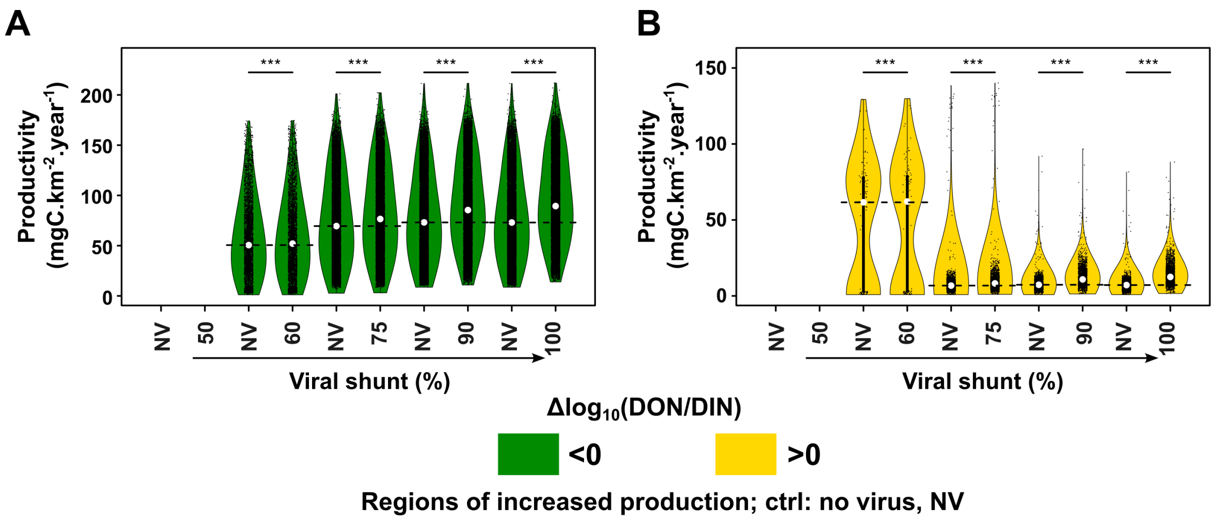


**Fig. S5. Impact of viral inclusion on the distribution of net primary productivity in regions of enhanced productivity relative to the simulation without the virus in function of the viral shunt.** Distribution of net primary productivity in regions of increased productivity relative to the no-virus simulation (NV), classified according to the sign of change in the $DNR$ ($log_{10}(DON/DIN)$, virus versus no virus) (**a**) $< 0$ and (**b**) $> 0$. For each viral shunt efficiency, comparison with the no-virus simulation is performed independently. Stars indicate paired Wilcoxon test significance ($p < 0.001$, Bonferroni correction for multiple testing). Note that for a viral shunt efficiency of 50%, a few grid points exhibited enhanced productivity, but they are not shown here because the sample size was too small to be considered representative ($n < 25$).

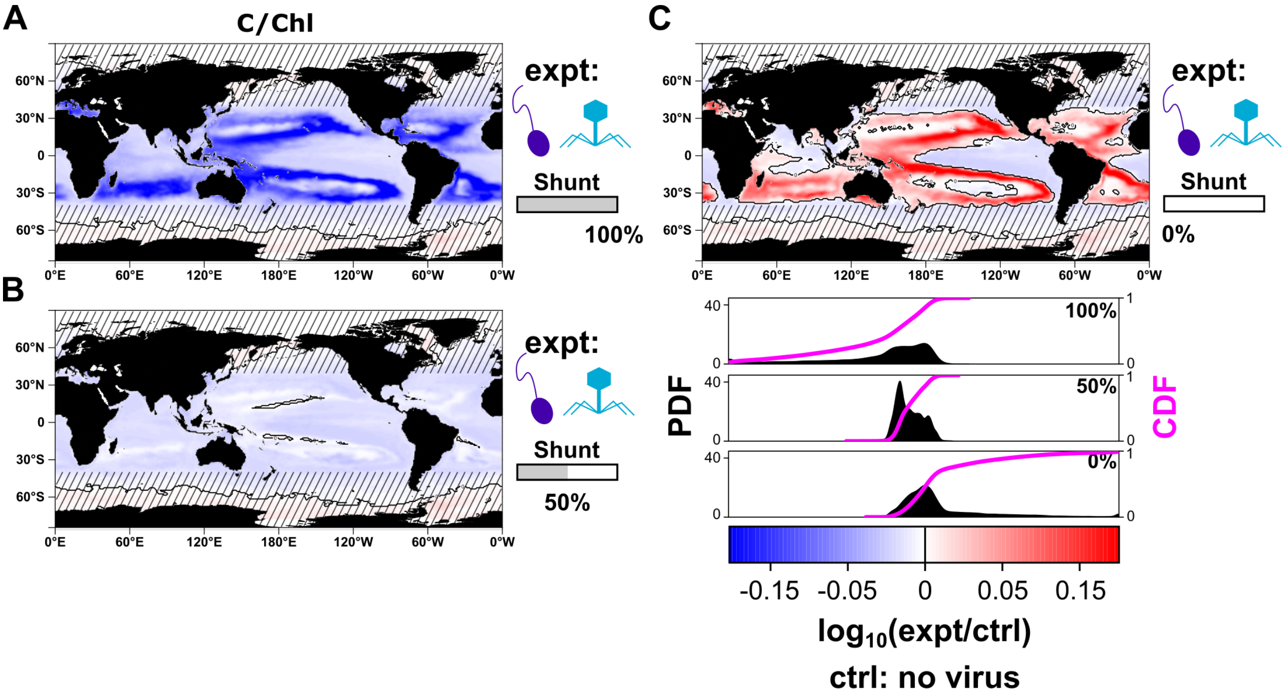


**Fig. S6. Impact of viral inclusion on the $C/Chl$ ratio in function of the viral shunt.** Time- and depth-averaged (one year, upper 100 m) differences in $C/Chl$ between the no virus simulation (control; ctrl) and simulation with the virus (expt) for shunt efficiencies of (**a**) 100%, (**b**) 75%, and (**c**) 50 %. Probability density function (PDF, black distribution) and cumulative distribution function (CDF, magenta) of fold changes for the different shunt efficiency values are shown in the bottom panels (data for the whole ocean). Absolute values of the $C/Chl$ ratio are shown in Fig. S3G-H for the simulation without viruses and the simulation with high viral shunt efficiency. Striped areas indicate regions outside the main niche of *Prochlorococcus* (poleward of 40°N/S). All $\log_{10}$ ratios are defined with the experiment (expt) simulation in the numerator and control (ctrl) simulation in the denominator. To avoid extreme values dominating the color scale, we capped absolute $log_{10}$ values at the 99th percentile of the distribution or at 2 (corresponding to a 100-fold change), whichever was smaller.

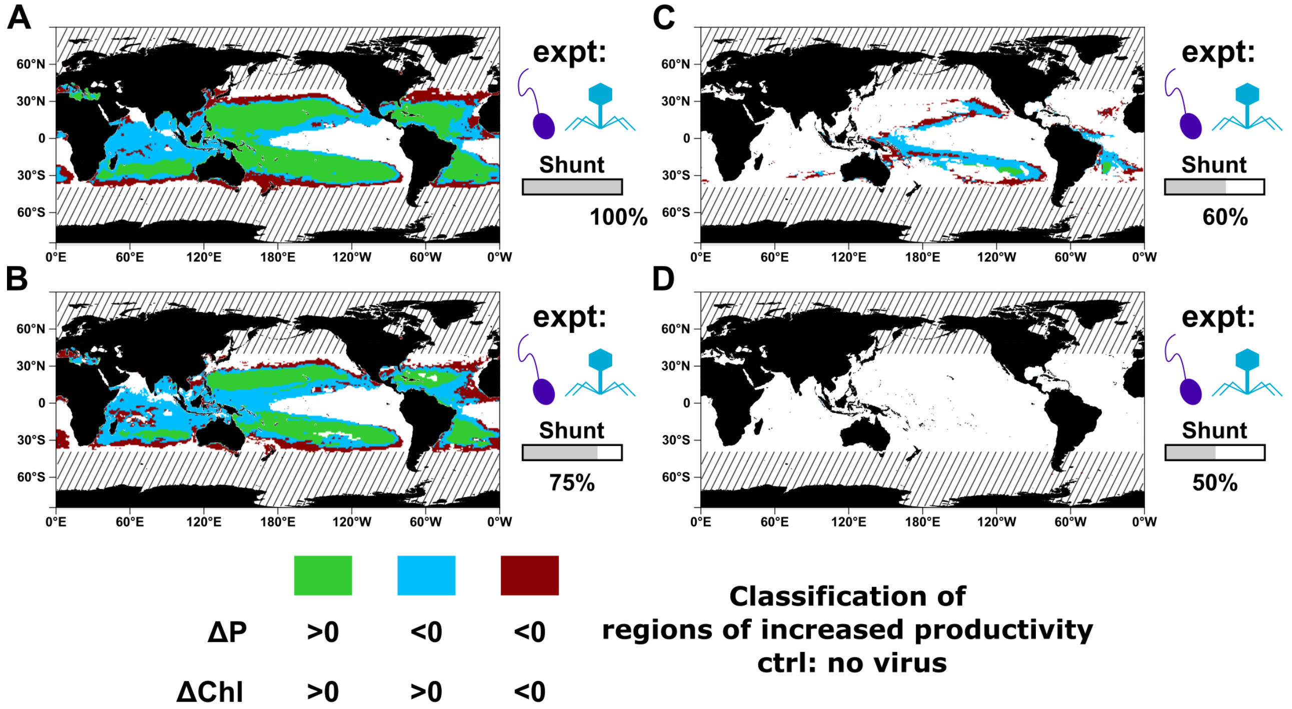


**Fig. S7. Classification of oceanic areas with increased productivity in function of the viral shunt based on changes in chlorophyll and carbon biomass relative to the no virus simulation.** (**a-d**) Regions of increased productivity are classified according to the signs of changes in time and depth averaged chlorophyll concentration (Chl) and P (total phytoplankton carbon concentration) relative to the averaged simulation without the virus (control; ctrl). Classification maps are shown for shunt efficiencies of (**e**) 100, (**f**) 75, (**g**) 60, and (**h**) 50%. For 50% shunt, almost no regions of increased productivity are found. Striped areas indicate regions outside the main niche of *Prochlorococcus* (poleward of 40°N/S).

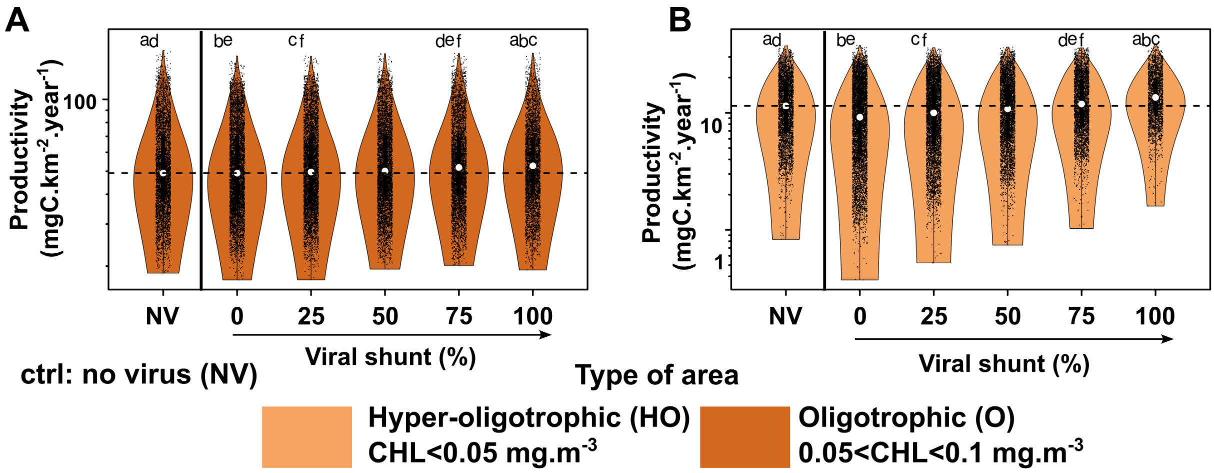


**Fig. S8. Effect of viral inclusion and the viral shunt efficiency on the productivity of oligotrophic regions.** Distribution of net primary productivity in (**a**) oligotrophic regions and (**b**) hyper-oligotrophic regions in function of the viral shunt and for the simulation without the virus. Letters indicate pairwise Wilcoxon test significance ($p < 0.05$, Bonferroni correction).

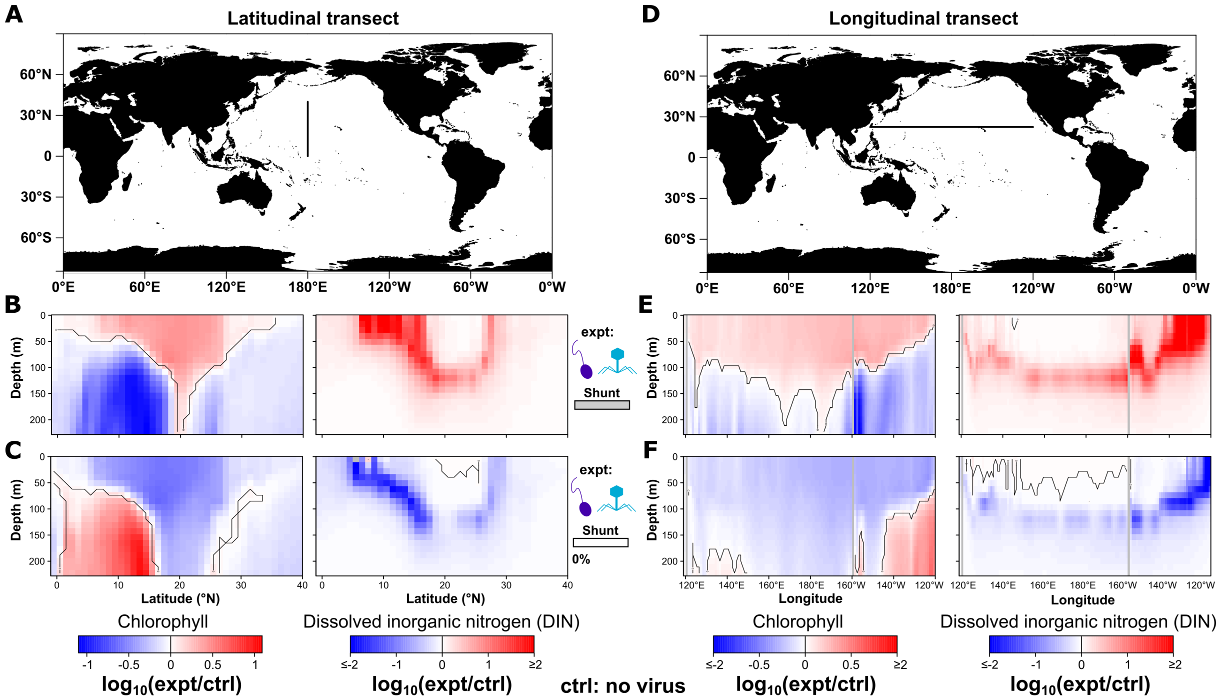


**Fig. S9. Regional changes in chlorophyll and total dissolved nitrogen in two transects of the North Pacific Subtropical Gyre in function of the viral shunt.** (**a**) Latitudinal transect (180°E). Changes in yearly averaged chlorophyll concentrations and total dissolved nitrogen compared to the simulation without viruses (control; ctrl) for viral shunt efficiencies of (**b**) 100, and (**c**) 50%. (**d**) Longitudinal transect (22.5°N). Changes in yearly averaged chlorophyll concentrations and total dissolved nitrogen compared to the simulation without viruses for viral shunt efficiencies of (**e**) 100, and (**f**) 50%. All $\log_{10}$ ratios are defined with the experiment (expt) simulation in the numerator and control (ctrl) simulation in the denominator. For the transect data presented here, we used the approx function from R to linearly interpolate values across an equally space depth grid of 20 points (between 0 and 220 m depth).
